# A conserved gene regulatory network controls root epidermal cell patterning in superrosid species

**DOI:** 10.1101/2023.02.20.529245

**Authors:** Yan Zhu, John Schiefelbein

## Abstract

- In superrosid species, root epidermal cells differentiate into root hair cells and non-hair cells. In some superrosids, the root hair cells and non-hair cells are distributed randomly (Type I pattern) and in others, they are arranged in a position-dependent manner (Type III pattern). The model plant Arabidopsis (*Arabidopsis thaliana*) adopts the Type III pattern, and the gene regulatory network (GRN) that controls this pattern has been defined. However, it is unclear whether the Type III pattern in other species is controlled by a similar GRN as in Arabidopsis, and it is not known how the different patterns evolved.
- In this study, we analyzed superrosid species *Rhodiola rosea*, *Boehmeria nivea,* and *Cucumis sativus* for their root epidermal cell patterns. Combining phylogenetics, transcriptomics, and cross-species complementation, we analyzed homologs of the Arabidopsis patterning genes from these species.
- We identified *R.rosea* and *B.nivea* as Type III species and *C.sativus* as Type I species. We discovered substantial similarities in structure, expression, and function of Arabidopsis patterning gene homologs in *R.rosea* and *B.nivea*, and major changes in *C.sativus*.
- We propose that in superrosids, diverse Type III species inherited the patterning GRN from a common ancestor, whereas Type I species arose by mutations in multiple lineages.

## Introduction

In multicellular organisms, cells adopt distinct identities during development to form appropriate spatial patterns. Since the establishment of Arabidopsis (*Arabidopsis thaliana*) as a model plant (Somerville & Koornneef, 2002), great progress has been made towards understanding pattern formation in this plant from embryogenesis to organogenesis (Capron *et al*., 2009; Causier *et al*., 2010; Cederholm *et al*., 2012; Shi & Vernoux, 2019; Manuela & Xu, 2020). However, the extent to which patterning mechanisms identified in Arabidopsis are also utilized by other plants is mostly unclear. In most vascular plants, root epidermal cells differentiate into two cell types: root hair cells which generate tip growing tubular-shaped structures called root hairs, and non-hair cells which do not generate root hairs. The distribution of root hair cells and non-hair cells within the epidermis (i.e., the root epidermal cell pattern) varies among different species.

In species adopting the Type I pattern, every root epidermal cell has the potential to differentiate into a root hair cell. This “random” pattern has been found in species across angiosperm groups, and has been proposed to represent the ancestral state (Dolan & Costa, 2001). In species adopting the Type II pattern, epidermal cells undergo a final asymmetric cell division, giving rise to a root hair cell and a non-hair cell. This pattern has been observed in extant basal angiosperms and monocots, but not in eudicots (Leavitt, 1904; Cutter & Feldman, 1970; Clowes, 2000; Kim & Dolan, 2011). In species adopting the Type III pattern, only epidermal cells located over the cleft of two underlying cortical cells differentiate into root hair cells. Accordingly, this is referred to as a position- dependent pattern. The cell position outside the cleft (anticlinal wall) of two cortical cells is defined as the Hair cell position (H-position), and the position outside a periclinal wall of a cortical cell is termed the Non-hair cell position (N- position). This Type III pattern was first identified in species belonging to the Brassicales order of the superrosid group (Cormack, 1935; Cormack, 1947; Bünning, 1951), and is now known to exist in many diverse eudicot species (Clowes, 2000; Pemberton *et al*., 2001) .

Arabidopsis, a member of the Brassicales order, exhibits the Type III root epidermal cell pattern (Dolan *et al*., 1993; Galway *et al*., 1994). Extensive molecular genetic analyses have led to the discovery of numerous patterning genes that act in a complex <u>g</u>ene regulatory <u>n</u>etwork (GRN). The downstream gene in this network, *<u>R</u>oot <u>H</u>air <u>D</u>efective <u>6</u>* (*RHD6*), is preferentially expressed in H-position cells and encodes a <u>b</u>asic <u>H</u>elix-<u>L</u>oop-<u>H</u>elix (bHLH) transcription factor that activates root hair development genes, leading to root hair outgrowth (Menand *et al*., 2007; Yi *et al*., 2010; Bruex *et al*., 2012). In N-position cells, *RHD6* transcription is suppressed by a <u>H</u>omeo<u>d</u>omain Leucine <u>Zip</u>per (HD-ZIP) family IV protein encoded by *<u>GL</u>ABRA*<u>2</u> (*GL2*) (Menand *et al*., 2007; Bruex *et al*., 2012; Lin *et al*., 2015). A transcription factor complex composed of R2R3 MYB protein <u>WER</u>EWOLF (WER), bHLH proteins <u>GL</u>ABRA<u>3</u> (GL3) or <u>E</u>nhancer of <u>GL</u>ABRA<u>3</u> (EGL3), and WD40-repeat protein <u>T</u>RANSPARENT <u>T</u>ESTA <u>G</u>LABRA<u>1</u> (TTG1) activates expression of *GL2* in N-position cells (Galway *et al*., 1994; Lee & Schiefelbein, 1999; Bernhardt *et al*., 2003; Song *et al*., 2011). The same <u>M</u>YB-<u>b</u>HLH-<u>W</u>D40 (MBW) complex also activates expression of *<u>C</u>A<u>P</u>RI<u>C</u>E* (*CPC*) in N-position cells. The R3 MYB protein encoded by *CPC* is able to move from the N-position cells to neighboring H- position cells through plasmodesmata and competes with WER for binding to bHLH partners, forming a CPC- GL3/EGL3-TTG1 complex that is unable to activate *GL2* (Wada *et al*., 1997; Wada *et al*., 2002; Koshino-Kimura *et al*., 2005; Kurata *et al*., 2005; Tominaga *et al*., 2007). *WER* is considered to play the critical role in patterning, as WER preferentially accumulates in N-position cells and directly binds to cis-regulatory elements of target genes, including *GL2* to determine non-hair cell fate, and *CPC* to determine root hair cell fate (Koshino-Kimura *et al*., 2005; Ryu *et al*., 2005). Upstream of this GRN is a putative positional signal, perceived by epidermal cells through the receptor-like-kinase encoded by *<u>SC</u>RA<u>M</u>BLED* (*SCM*) (Kwak *et al*., 2005), leading to differences in *WER* expression in N-position cells and H- position cells (Kwak & Schiefelbein, 2007).

Homologs of the Arabidopsis root epidermis patterning genes have been identified from diverse land plants (Kim & Dolan, 2016; Chopra *et al*., 2019; Zhang & Hülskamp, 2019; Cai *et al*., 2020). However, they have not been investigated from other Type III species outside the Brassicales order. Therefore, it is unclear whether the molecular mechanism controlling root epidermal cell patterning in Arabidopsis is conserved in diverse Type III species. Here, we describe the identification and analysis of two superrosid species outside the Brassicales order that adopt the Type III root epidermal cell pattern. The results indicate that these species use the same molecular mechanism as Arabidopsis for their root epidermal patterning, suggesting that this regulatory mechanism was likely inherited from a common ancestor. We also discovered abnormalities in the structure and/or expression of these patterning gene homologs in Type I species, implying that Type I species arise from mutations in the Type III GRN. Together, our study provides insights into the evolution of cell type patterning in plants and extends our knowledge of the molecular basis of root epidermal cell fate beyond the model species Arabidopsis.

## Material and Methods

### Plant materials and growth conditions

*Arabidopsis thaliana* seeds were surface sterilized and grown as previously described (Estelle & Somerville, 1987). Arabidopsis lines used in this study have been reported previously: *ProAtGL2:GUS* (Masucci *et al*., 1996), *rhd6-1* (Masucci & Schiefelbein, 1994), *gl2-1* (Koornneef, 1981), *wer-1 ProAtGL2:GUS* (Lee & Schiefelbein, 1999), and *cpc-1 ProAtGL2:GUS* (Lee & Schiefelbein, 2002).

*Boehmeria nivea* seeds (Ebay) were surface sterilized with bleach solution (30% (v/v) bleach plus 0.02% (v/v) Triton X-for 10 min and sown on Gelrite-solidified nutrient plates (for root hair counting assay) or on nylon mesh topped agar- solidified 0.5× Murashige and Skoog (MS) medium (pH 5.7- 5.8) with 1% sucrose (for root zonation RNA sample collection), then stratified at 4°C for 3 days. *Rhodiola rosea* seeds (Strictly Medicinal (https://strictlymedicinalseeds.com/)) were surface sterilized with bleach solution for 10 min, washed with water for 5 times, then soaked in 50 µg/L GA3 and stratified at 4°C for 1 week. The imbibed seeds were washed with water once, surface sterilized with bleach solution again for 10 min, and washed with water three times. The seeds were sown on Gelrite-solidified nutrient media (topped with nylon mesh for root zonation RNA sample collection). Both *B.nivea* and *R.rosea* seedlings were grown in a chamber set to 22°C with continuous light.

*Cucumis sativus* ’Marketmore’ seeds (Strictly Medicinal (https://strictlymedicinalseeds.com/)) were surface sterilized with bleach solution for 10 min, washed with water for 5 times, then soaked in 50 µg L^-1^ GA3 and stratified at 4°C for 2 days. The imbibed seeds were washed with water once, surface sterilized with 15% hydrogen peroxide for 15 min, then washed with water three times. The seeds were sown on Gelrite-solidified mineral nutrient media and grown in a chamber set to 22°C with continuous light.

### Microscopy

Before being subjected to microscopy analysis, Arabidopsis seedlings were grown for 4-5 days and *R.rosea*, *B.nivea*, and *C.sativus* seedling roots were grown to 2-3 cm.

The root hair counting assay was essentially performed as previously described (Bernhardt *et al*., 2003) (**Methods S1**). Raw root hair counting data is included as **Dataset 1**.

Analysis of the transgenic plants bearing a GUS reporter gene was performed as previously described (Masucci *et al*., 1996).

Plastic transverse sections were obtained as previous described (Masucci *et al*., 1996) with modifications, and detailed procedures are described in **Methods S2**.

The ClearSee protocol (Kurihara *et al*., 2015; Ursache *et al*., 2018) was used for fluorescence signal detection with a Leica SP5 laser scanning confocal microscope.

To observe trichome phenotypes, Arabidopsis seedlings were grown in the soil and the 3^rd^ and 4^th^ true leaves were observed and photographed under a stereo microscope.

The seed coat mucilage was analyzed by imbibing seeds in water for 5 min, staining with 0.01% (w/v) ruthenium red for 1 h, then washing with water twice before observation under a stereo microscope.

### Transgene constructs and plant transformation

Oligonucleotides used in this study are listed in **Dataset 2**. All plasmids were constructed using the binary vector *pCAM-EYFP-C1* (Preuss *et al*., 2004), a kind gift from Erik Nielsen (University of Michigan). In brief, to generate the constructs for complementation experiments, the *CaMV35S* promoter and *EYFP* cDNA sequences (CDS) were replaced by promoters, *GFP*/*mCherry* CDSs, and 3’ untranslated regions (UTRs) of interest to generate backbones, then the CDSs of genes were ligated or assembled in frame with the *GFP*/*mCherry* CDS. For constructs expressing *CPL*s, the *HygR* gene was replaced by a *bar* gene cloned from the binary vector *pCB302* (Xiang *et al*., 1999). To generate the promoter- reporter constructs, the *CaMV35S* promoter and *EYFP* CDS were replaced by sequences encoding the signal peptide of *AtWAK2* (*Sp^WAK2^*), an in frame *mCherry* CDS, and a sequence encoding an ER retention signal HDEL. The promoter regions of interest were then inserted upstream of the *Sp^WAK2^- mCherry-HDEL* cassette. Detailed procedures are described in **Methods S3**.

All constructs were verified by sequencing and transferred into *Agrobacterium tumefaciens* strain GV3101. Arabidopsis seedlings were transformed through floral dipping as described (Clough & Bent, 1998). The T1 seedlings of *CPL* related lines were screened for glufosinate ammonium (50 mg L^-1^ in growth media) resistance. T1 seedlings of all other lines were screened for hygromycin (50 mg L^-1^ in growth media) resistance. High efficiency TAIL-PCR (Liu & Chen, 2007) was used to map insertion sites of *ProAtGL2:GFP-CsGLH1* transgenes.

### RNA isolation and sequencing

For *R.rosea* and *B.nivea*, root zonation RNA samples were collected from 2-3 cm long primary roots. The root developmental zones were defined as previously described (Huang & Schiefelbein, 2015) (**Fig. S1**). Non-stranded library construction and sequencing were performed by the University of Michigan Advanced Genomics Core (**Methods S4)**.

### RNA-seq analysis

Published genome and transcriptome information and NCBI Sequence Read Archive accession numbers for published sequencing data used in this study are listed in **Dataset 3**.

*A. thaliana*, *B.nivea*, and *C.sativus* reads were processed as previously described (Huang & Schiefelbein, 2015) and mapped to corresponding genomes using STAR (Version 2.7.4a) (Dobin *et al*., 2013) with 2-pass mode followed by quantification using RSEM (Version 1.3.3) (Li & Dewey, 2011). *R.rosea* reads were filtered for ribosomal RNA, corrected for erroneous kmers and trimmed before being assembled using Trinity (Grabherr *et al*., 2011) and rnaSPAdes (Version 3.15.5) (Bushmanova *et al*., 2019). The outputs were concatenated using EvidentialGene (Gilbert, 2019) to generate the final assembly. The processed reads were mapped back to the assembly and quantified using RSEM (Version 1.3.3) (Li & Dewey, 2011). The *R.rosea* transcriptome was annotated using Trinotate pipeline (Version 3.2.2) (Bryant *et al*., 2017).

Details on reads processing, mapping, *de novo* assembling, and annotating can be found in **Methods S5**.

Differential expression analysis was performed using edgeR (Version 3.40.0) (Robinson *et al*., 2010) (**Methods S5**). Genes with a |log2(fold-change)| ≥ 1 and a false discovery rate ≤ 0.05 between the compared developmental zones were retained and present in **Dataset 4**.

### Phylogenetic analysis

Information of genome and transcriptome annotations used and accession numbers of protein coding genes are listed in **Dataset 3**. The pipeline was modified from one previously described (Huang & Schiefelbein, 2015) (**Methods S6**).

### Synteny network analysis

All the genome information used is listed in **Dataset 3**. The SynNet pipeline (Zhao *et al*., 2017; Zhao & Schranz, 2019) was used to construct microsynteny networks of genes of interest with the following setting: diamond blast -k 5, MCScanX -s 5 -m 25. The resulting synteny network was visualized by Cytoscape (Version 3.9.1) (Shannon *et al*., 2003).

## Results

### Root epidermal cell patterns in superrosid species

We surveyed multiple superrosid species with a particular focus on species with reported Type I or Type III patterns or species with available genome/root transcriptome resources. For this study, we conducted detailed morphological and molecular analyses on *Boehmeria nivea*, *Cucumis sativus*, and *Rhodiola rosea*, which are superrosid species distantly related to Arabidopsis (**Fig. 1a**).

**Fig. 1.**
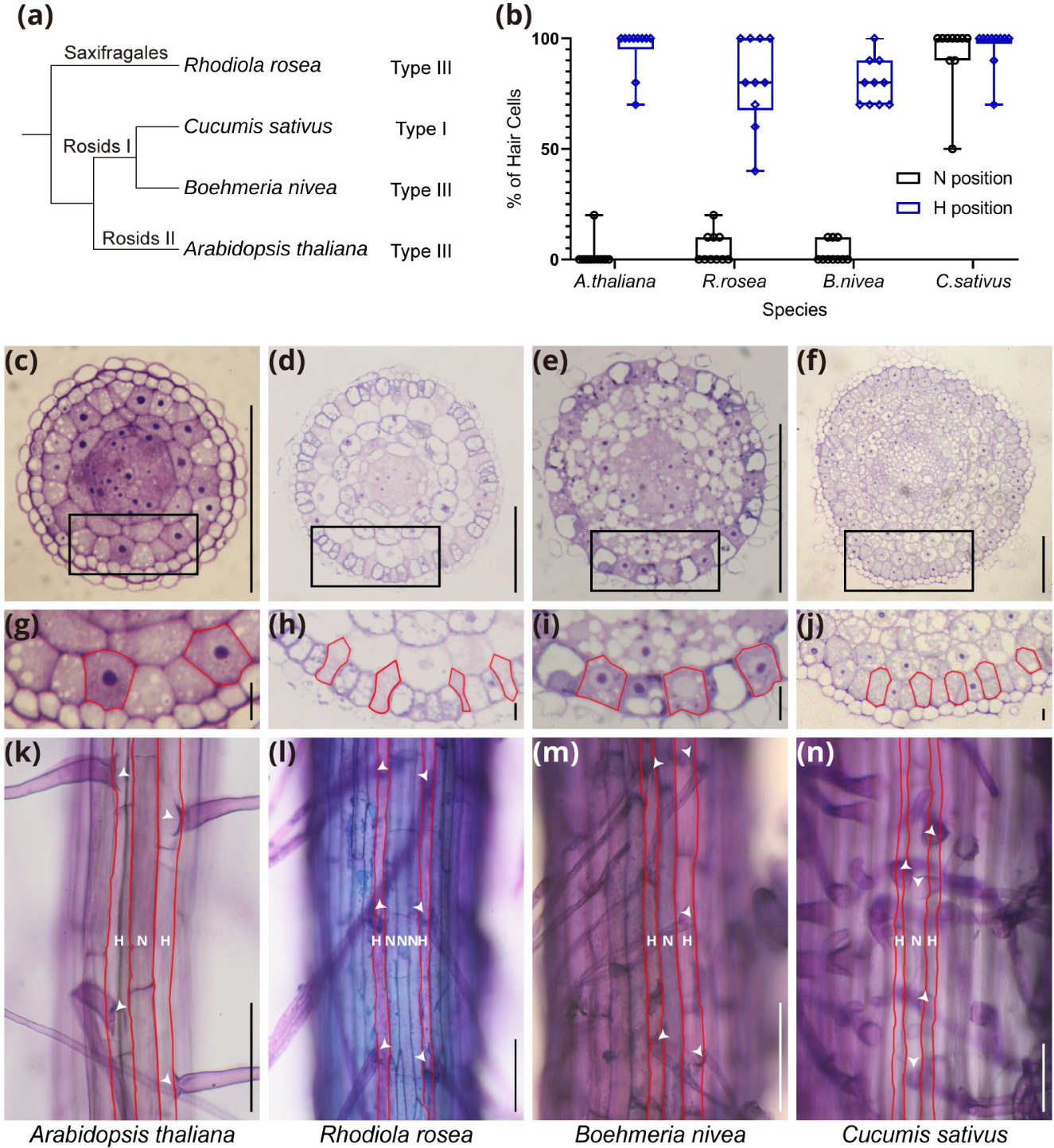
Root epidermal cell patterns of selected species. (**a**) A simplified cladogram of selected species and root epidermal cell patterns adopted by the species, respectively. (**b**) Root epidermal patterns of selected species quantified by root hair counting assay. (**c**-**f**) Transverse sections of the primary root tips of selected species stained with toluidine blue. Bar=100um. (**g**-**j**) Zoom in view of the regions labelled in (**c**-**f**). H-position cells are outlined by red lines. Bar=10um. (**k**-**n**) Primary roots of selected species stained with toluidine blue. N-position cells are labelled by N, and H-position cells are labelled by H. H-position cell files are also outlined by red lines. Arrowheads point to root hairs. Bar=100um.

First, the pattern of root epidermal cells in primary seedling roots of these species was analyzed. *A.thaliana*, *R.rosea,* and *B.nivea* were found to exhibit the Type III pattern, whereas *C.sativus* roots showed the Type I pattern (**Fig. 1a**).

Prior to root hair formation, the H-position cells of Type III species are known to exhibit more intense cytoplasmic staining and reduced vacuolation relative to the N-position cells (Cormack, 1935; Bünning, 1951; Dolan *et al*., 1993). We confirmed greater relative cytosolic density in H-position cells of *A.thaliana*, *R.rosea*, and *B.nivea*, but not *C.sativus*. Further, we observed differential vacuolation between H- and N- position cells in the root sections from *A.thaliana*, *R.rosea*, and *B.nivea*. (**Fig. 1c-j**). These observations show that roots of *R.rosea* and *B.nivea* have the typical features of Type III species (like *A.thaliana*), whereas roots of *C.sativus* show Type I features.

We also found that H-position cells of *R.rosea* stained purple and N-position cells stained blue (**Fig. 1l**). This distinction was also observed in the *B.nivea* root elongation region (**Fig. S1**), but it is not observed in roots of *A.thaliana* or *C.sativus* (**Fig. 1k,n**). These results suggest that the *R.rosea* and *B.nivea* may share additional H-position and N-position cell differences not present in *A.thaliana*.

### Identification and functional analysis of *RHD6* homologs

In Arabidopsis, *RHD6* is the key transcriptional regulator of root hair initiation and the downstream target of the root epidermal patterning GRN (Huang *et al*., 2017). Therefore, we sought to investigate the conservation of *RHD6* and its regulation in our selected species.

We first identified RHD6-like bHLH (RLB) proteins from genomes or transcriptomes of these species (**Dataset 3**). *R.rosea*’s sister species, *Rhodiola crenulata*, and a basal eudicot species *Trochodendron aralioides* were included in this analysis to better evaluate orthology of the RLBs. Maximum likelihood analysis resolved a monophyletic group containing AtRHD6 and AtRSL1, 2 RrRLBs, 2RcRLBs, 1 BnRLB, 1 CsRLB, and 3 TaRLBs (**Fig. 2a**). Amino acid sequence alignment of these RLBs demonstrated conservation of the bHLH domain and the RSL domain (Kim & Dolan, 2016) (**Fig. S2**). The available genome information (from At, Bn, Rc, Cs, and Ta) allowed us to perform synteny network analysis (Zhao *et al*., 2017; Zhao & Schranz, 2019), which showed that *AtRHD6*, *AtRSL1*, *BnRLB*, and the 3 *TaRLBs* are in a microsynteny network (**Fig. S3a**).

**Fig. 2.**
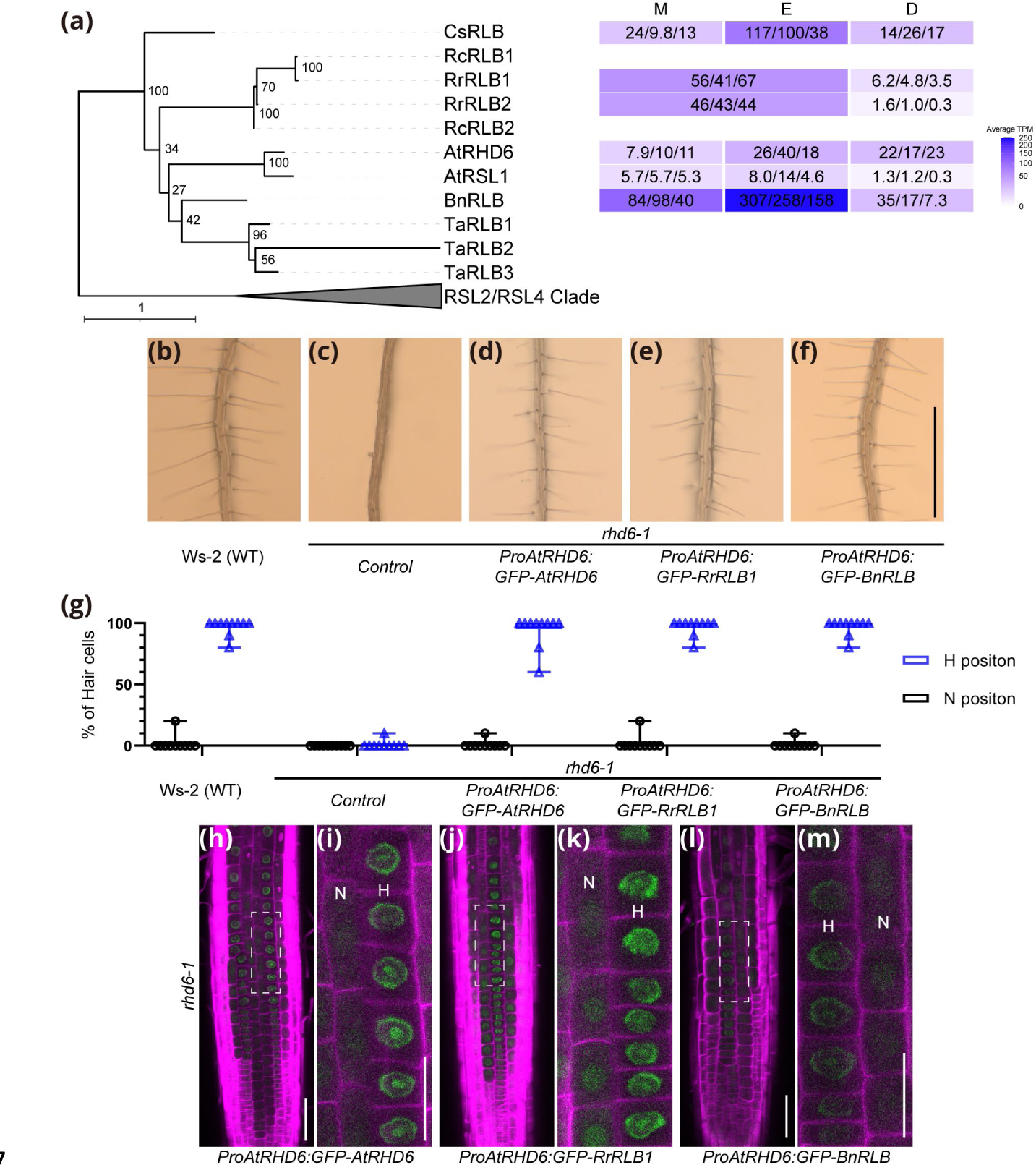
Identification and functional analysis of *RLB*s. (**a**) Left, maximum likelihood tree of RLB proteins. The tree is rooted to the RSL2/4 clade. The scale bar refers to amino acid substitutions per site. Right, gene expression levels in root developmental zones (<u>M</u>eristem, <u>E</u>longation, <u>D</u>ifferentiation). TPM values of 3 RNA-seq replicates are shown in each box, and the average values are visualized by heatmaps after square root transformation. (**b**-**f**) Root hair phenotypes of Arabidopsis *WS-2*(WT), *rhd6-1*, and *rhd6-1* mutant seedlings bearing *ProAtRHD6:GFP-RLB* constructs. All transgenic lines shown are representative T3 individuals. Bar=1mm. (**g**) Quantification of root epidermal cell phenotypes of the lines shown in (**b**-**f**). Root hair counting raw data and information of other independent lines are listed in **Dataset 1**. (**h**, **j**, **l**) Confocal images of roots from the lines shown in (**d**-**f**). Bar= 50um. (**i**, **k**, **m**) Zoom in view of the regions highlighted by dash lines in (**h**, **j**, **l**), respectively. N-position cell files are labelled by N, and H-position cell files are labelled by H. Bar=25um. GFP signals are pseudocolored as green, CalcoFluor White signals are pseudocolored as magenta.

To examine expression patterns of the identified *RLBs* in the root, RNA from the root developmental zones (meristem (M), elongation (E), and differentiation (D)) of *R.rosea* and *B.nivea* roots (**Fig. S1b,c**) were isolated, sequenced and compared to previously obtained RNA-seq data from the same zones of *A.thaliana* and *C.sativus* (Huang & Schiefelbein, 2015). *AtRHD6* and *AtRSL1* transcripts were detected throughout the three developmental zones with *AtRHD6* transcripts preferentially accumulating in elongation and differentiation zones of the root (**Fig. 2a**; **Dataset 4, At_MvsD, At_MvsE)** and *AtRSL1* transcripts preferentially accumulating in meristem and elongation zones (**Fig. 2a**; **Dataset 4, At_MvsD, At_EvsD)**. Transcripts of *RrRLB1*, *RrRLB2*, *BnRLB*, and *CsRLB* were detected in all 3 developmental zones of the roots, with *RrRLBs* preferentially expressed in the meristem/elongation zone (**Fig.2a**; **Dataset 4, Rr_MEvsD),** and *BnRLB* and *CsRLB* preferentially expressed in the elongation zone (**Fig. 2a; Dataset 4, Bn_MvsE, Bn_EvsD, Cs_MvsE, Cs_EvsD**).

We next sought to examine whether *RLB*s have conserved function. The homologs with greatest root transcript accumulation, *RrRLB1* and *BnRLB*, and the *AtRHD6* control, were selected for molecular complementation experiments to produce GFP-RLB fusion proteins under the control of the *AtRHD6* promoter (*ProAtRHD6*) in *rhd6-1* mutant plants. Compared to the Wassilewskija-2 (WS-2) wild type (WT) roots, *rhd6-1* mutant roots were void of root hairs (**Fig. 2b,c,g**), and the *ProAtRHD6:GFP-AtRHD6* construct effectively rescued this hairless phenotype (**Fig. 2d,g**). Further, the GFP-AtRHD6 fusion proteins accumulated preferentially in H-position cells, although weak GFP signals could also be observed in certain N-position cells (**Fig. 2h,i**). The GFP signal was found in periphery of the nucleus and in the nucleolus in H-position cells (**Fig. 2h,i**). Both the *ProAtRHD6:GFP-RrRLB1* and the *ProAtRHD6:GFP-BnRLB* constructs could rescue the hairless phenotype of *rhd6-1* mutant plants (**Fig. 2e-g; Dataset 1**). In addition, the GFP-RrRLB1 and GFP-BnRLB fusion proteins exhibited preferential accumulation and subcellular localization patterns similar to GFP-AtRHD6 (**Fig. 2j-m**). These results suggest that RLB proteins from different Type III species possess conserved functions in promoting root hair formation.

### Identification and functional analysis of *GL2* homologs

In Arabidopsis, *AtRHD6* expression is suppressed in N- position cells by the GL2 protein (AtGL2), causing these cells to differentiate into non-hair cells (Masucci *et al*., 1996).

<u>G</u>L2-like <u>H</u>D-Zip (GLH) proteins were identified from each of the selected species (**Fig. 3a**). The proteins showed conservation of the Homeodomain and the START domain (**Fig. S4**) and the *AtGL2*, *RcGLH1*, *BnGLH*, *CsGLH1*, and *TaGLH* genes were found to have shared synteny (**Fig. S3b**). In Arabidopsis roots, *AtGL2* transcripts accumulate throughout all three root developmental zones and *GLHs* from other Type III species (*BnGLH* and *RrGLH1*) showed similar expression profiles. The *C.sativus GLH*s (*CsGLH1* and *CsGLH2*) showed low transcript level (**Fig. 3a; Dataset 4**). This suggests a possible linkage between *GLH* expression and the Type III root epidermal cell pattern.

**Fig. 3.**
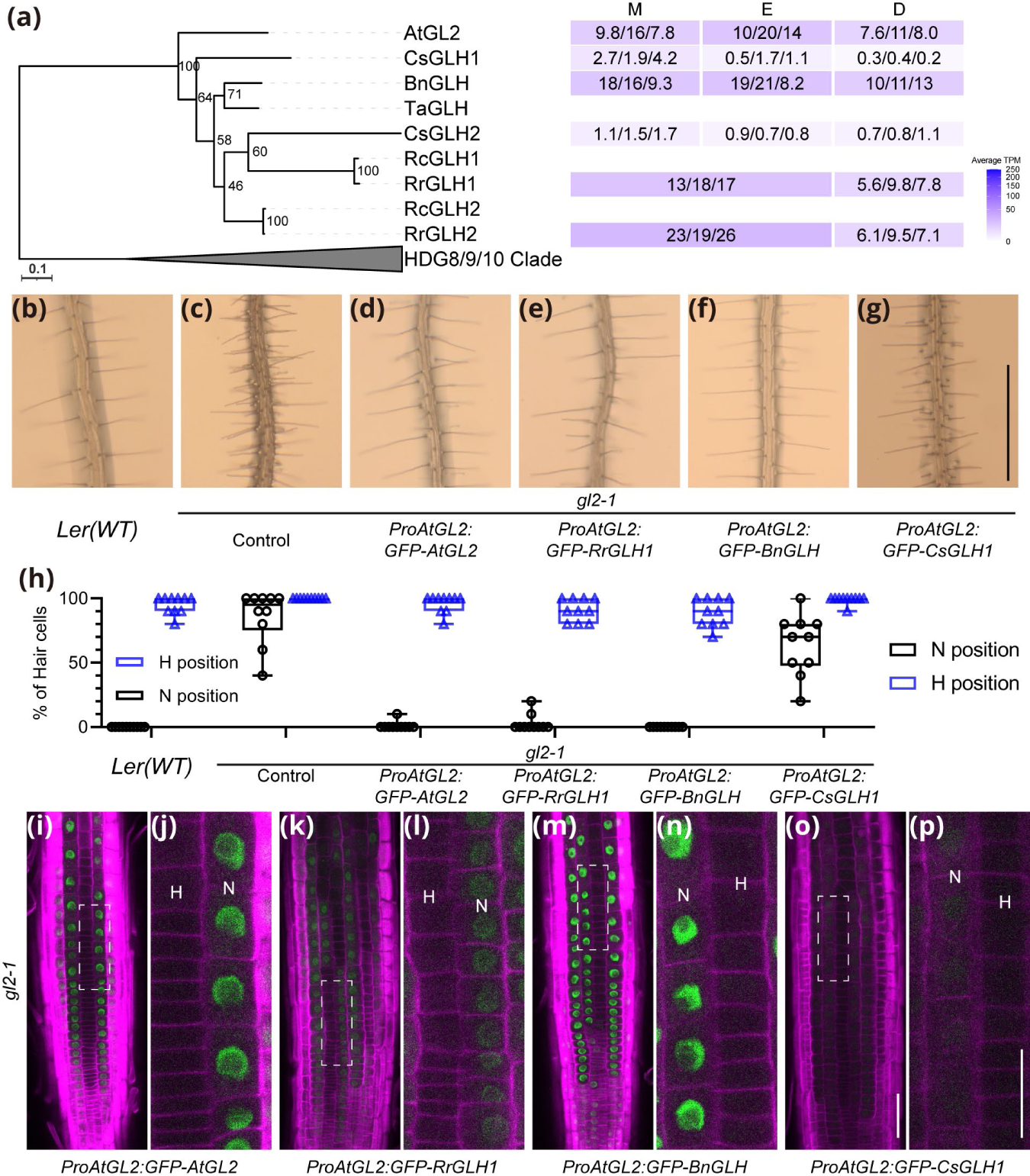
Identification and functional analysis of *GLH*s. (**a**) Left, maximum likelihood tree of GLH proteins. The tree is rooted the HDG8/9/10 clade. The scale bar refers to amino acid substitutions per site. Right, gene expression levels in root developmental zones. TPM values of 3 RNA-seq replicates are shown in each box, and the average values are visualized by heatmaps after square root transformation. (**b**-**g**) Root hair phenotypes of Arabidopsis *Ler*(WT), *gl2-1* and *gl2- 1* mutant seedlings bearing *ProAtGL2:GFP-GLH* constructs. All transgenic lines shown are representative T3 individuals. Bar=1mm. (**h**) Quantification of root epidermal cell phenotypes of the lines shown in (**b**-**g**). Root hair counting raw data and information of other independent lines are listed in **Dataset 1**. (**i**, **k**, **m**, **o**) Confocal images of roots from the lines shown in (**d**-**g**). Bar=50um. (**j**, **l**, **n**, **p**) Zoom in view of the regions highlighted by dash lines in (**i**, **k**, **m**, **o**). N-position cell files are labelled by N, and H-position cell files are labelled by H. Bar=25um. GFP signals are pseudocolored as green, CalcoFluor White signals are pseudocolored as magenta.

To investigate regulation of *GL2* homolog expression, we fused the cis-regulatory region of *BnGLH* (*ProBnGLH*) to the ER-localized marker gene *Sp^WAK2^-mCherry-HDEL* (Nelson *et al*., 2007) and introduced this construct into Arabidopsis WT plants. In roots of the resulting transgenic seedlings, mCherry signals could be observed in N-position cells (**Fig. S5a**), suggesting conservation of cis-regulatory elements of the *B.nivea GL2* homolog.

We next sought to examine whether *GLH*s have conserved functions. Based on their expression pattern and synteny, *RrGLH1*, *BnGLH*, *CsGLH1*, and *AtGL2*, were selected for molecular complementation experiments. As expected, the control *ProAtGL2:GFP-AtGL2* construct rescues the excessive root hair phenotype of Arabidopsis *gl2-1* mutant roots (**Fig. 3b-d,h**), and the GFP-AtGL2 fusion protein was found to preferentially accumulate in nuclei of N-position cells (**Fig. 3i-j**). The *ProAtGL2:GFP-RrGLH1* and *ProAtGL2:GFP- BnGLH* constructs could also rescue the excessive root hair phenotype of *gl2-1* mutant plants (**Fig. 3e,f,h; Dataset 1**), and the fusion proteins had similar accumulation pattern as GFP- AtGL2 (**Fig. 3k-n**). A *gl2-1* mutant line potentially homozygous for a single *ProAtGL2:GFP-CsGLH1* transgenic insertion exhibited partial rescue (**Fig. 3g-h, Dataset 1**) and consistent with this phenotype, weak fluorescent signals of GFP- CsGLH1 fusion protein were observed in N-position cells. (**Fig. 3o-p**). We also analyzed a population containing multiple *ProAtGL2:GFP-CsGLH1* transgenic insertions, and found seedlings showing the rescued phenotype comparable to those expressing *GLH*s from Type III species (**Fig. S6a-d; Dataset 1**). Together, these results indicate that GL2 homolog proteins from these species possess conserved function in mediating the Type III pattern.

In Arabidopsis, *GL2* also regulates trichome formation (Rerie *et al*., 1994) and seed coat mucilage synthesis (Western *et al*., 2001). *ProAtGL2:GFP-AtGL2*, *ProAtGL2:GFP-RrGLH1*, *ProAtGL2:GFP-BnGLH*, and multiple copies of *ProAtGL2:GFP-CsGLH1* were able to restore trichome formation on the adaxial side of true leaves (**Figs. S7, S6e)**, as well as the seed mucilage accumulation (**Figs. S8, S6f**) of the *gl2-1* mutant. These results suggest general conservation of all AtGL2 protein functions for these GLHs when appropriately produced in the various Arabidopsis tissues.

### Identification of *TTG1* and *GL3*/*EGL3* homologs

In the developing Arabidopsis root epidermis, *AtGL2* expression is activated by an MBW complex (WER- GL3/EGL3-TTG1) in N-position cells, which specifies the non- hair cell fate.

<u>T</u>TG1-like <u>W</u>D40 (TLW) proteins from the selected species were used to build a ML tree revealing a monophyletic clade containing AtTTG1, RcTLW, RrTLW, BnTLW, CsTLW, and 2 TaTLWs (**Fig. S9a**). Alignment of amino acid sequences demonstrated conservation of the four WD40 repeats and the phosphorylation site (Li *et al*., 2018) of AtTTG1 (**Fig. S10**). Synteny analysis suggested that *AtTTG1*, *RcTLW*, *BnTLW*, *TaTLW1*, and *TaTLW2* genes had shared synteny (**Fig. S9c**). In Arabidopsis, *AtTTG1* is expressed throughout all 3 developmental zones of the root. *RrTLW* and *BnTLW* showed similar expression profiles as *AtTTG1*. *CsTLW* was also expressed throughout the root, but lower transcript abundance (**Fig. S9a**).

We identified <u>G</u>L3-like <u>b</u>HLH (GLB) proteins from Type III species, which included a monophyletic clade containing AtGL3, AtEGL3, BnGLB, and 4 RcGLBs and 4 RrGLBs (**Fig. S9b**). However, no GLBs were identified from *C.sativus* or *T.aralioides*.

We also identified a sister clade to the GLB clade, which contains <u>M</u>YC1-like <u>b</u>HLH (MLB) proteins. Two *Rhodiola rosea* <u>M</u>YC1/<u>G</u>L3 like <u>b</u>HLH (RrMGB) proteins were found clustered with the whole MLB/GLB clade (**Fig. S9b**). Some of these proteins lack certain domains when compared to AtGL3, while the remaining ones shared conserved <u>J</u>az-Interacting <u>D</u>omain (JID), <u>T</u>ranscription <u>A</u>ctivation <u>D</u>omain (TAD), bHLH domain, and <u>C</u>-terminal <u>D</u>omain (CD), as well as the residues required for R2R3 MYB interaction (Wang *et al*., 2021) (**Figs. S11,S12**). Interestingly, our synteny analysis revealed a network including both *GLB*s and *RLB*s (**Fig. S9d**), suggesting a common ancestry of these genes.

In Arabidopsis, *GL3* and *EGL3* are expressed preferentially in the meristem and elongation zones (**Fig. S9b**; **Dataset 4, At_MvsD, At_EvsD**). Transcripts of *RrGLB2*, which encodes the only full length RrGLB, also preferentially accumulated in the meristem/elongation zone (**Fig. S9b; Dataset 4, Rr_MEvsD**). *BnGLB* transcripts were also detected in the roots (**Fig. S9b**). Together, these results indicate conservation of *GL3* homologs in species adopting the Type III pattern, whereas the lack of a *GLB* in *C.sativus* may contribute to its Type I pattern.

### Identification and functional analysis of *WER* homologs

Previous studies have demonstrated the R2R3 MYB component in the MBW complex is trait specific (Tian & Wang, 2020). The trait-specific MYB protein for Arabidopsis Type III root hair pattern is WER, so we next focused on analyzing its homologs in our selected species.

We were able to identify <u>W</u>E<u>R</u>-like (WRL) MYB proteins from *A.thaliana*, *R.rosea*, *B.nivea*, *R.crenulata* and the basal eudicot *T*.*aralioides* (**Fig. 4a**). Interestingly, we were not able to identify any WRL from the Type I species *C.sativus*. Amino acid sequence alignment suggested that all WRLs are similar in length and presence of canonical R2 and R3 MYB repeats, except for RrWRL1 and RcWRL2 which lack part of the R2 MYB repeat, and TaWRL2 which has an extended C terminus (**Fig. S13**). The alignment also revealed conservation of key residues previously shown to be required for bHLH protein interaction and DNA sequence specific recognition (Wang *et al*., 2019; Wang *et al*., 2021) among the WRLs (**Fig. S13**). Synteny analysis suggested that all *WRL*s identified from *A.thaliana*, *B.nivea*, *R.crenulata*, and *T. aralioides* had shared synteny except *TaWRL4* (**Fig. S3c**), indicating a common ancestry.

**Fig. 4.**
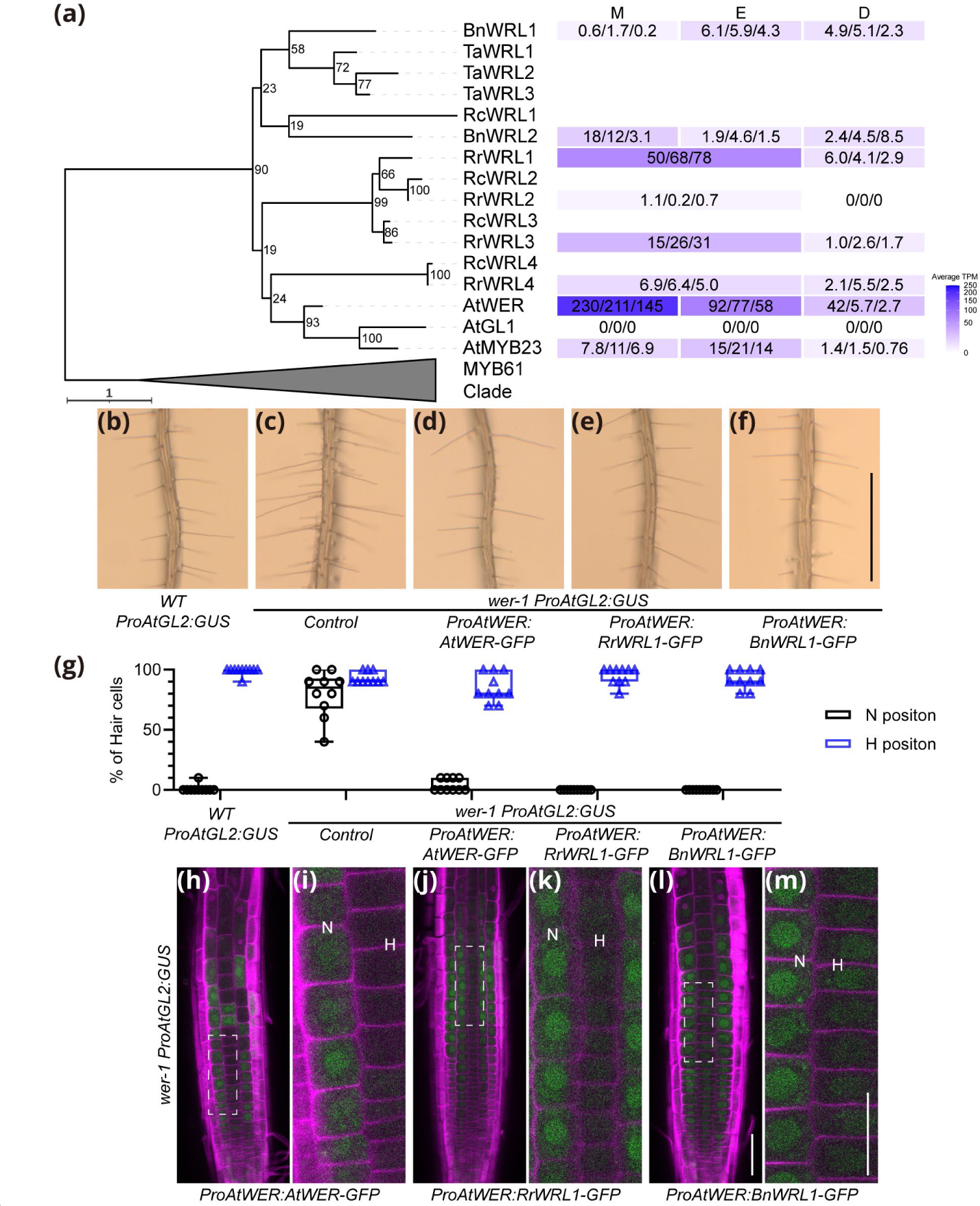
Identification and functional analysis of *WRL*s. (**a**) Left, maximum likelihood tree of WRL proteins. The tree is rooted the MYB61 clade. The scale bar refers to amino acid substitutions per site. Right, gene expression levels in root developmental zones. TPM values of 3 RNA-seq replicates are shown in each box, and the average values are visualized by heatmaps after square root transformation. (**b**-**f**) Root hair phenotypes of Arabidopsis WT, *wer-1* and *wer-1* mutant bearing *ProAtWER:WRL-GFP* constructs. Each line also carries a *ProAtGL2:GUS* transgene. All transgenic lines shown are representative T3 individuals. Bar=1mm. (**g**) Quantification of root epidermal cell phenotypes the lines shown in (**b**-**f**). Root hair counting raw data and information of other independent lines are listed in **Dataset 1**. (**h**, **j**, **l**) Confocal images of roots from the lines shown in (**d**-**f**). Bar=50um. (**i**, **l**, **m**) Zoom in view of the regions highlighted by dash lines in (**h**, **j**, **l**). N-position cell files are labelled by N, and H-position cell files are labelled by H. Bar=25um. GFP signals are pseudocolored as green, CalcoFluor White signals are pseudocolored as magenta.

In Arabidopsis roots, *AtWER* is preferentially expressed in the meristem and elongation zones, *AtMYB23* is preferentially expressed in the meristem and elongation zones at a lower level compared to *AtWER*, and *AtGL1* is not expressed (**Fig. 4a**; **Dataset 4, At_MvsD, At_EvsD**). *RrWRL1* and *RrWRL3* transcripts preferentially accumulated in the meristem/elongation zone of the root, whereas *RrWRL4* transcripts were detected throughout all three zones (**Fig. 4a**; **Dataset 4, Rr_MEvsD**). Transcripts of *BnWRL1* preferentially accumulated in the elongation and differentiation zones and transcripts of *BnWRL2* were detected throughout all 3 root developmental zones (**Fig. 4a**; **Dataset 4, Bn_MvsE, Bn_MvsD**).

At the cellular level, *AtWER* was reported to be expressed in both N- and H-position cells, with stronger expression in N- position cells (Lee & Schiefelbein, 2002). This expression pattern was further confirmed by the *ProAtWER:Sp^WAK2^- mCherry-HDEL* construct introduced into WT Arabidopsis plants (**Fig. S5b**). We generated corresponding reporter constructs using the cis-regulatory sequences of the *BnWRL*s fused to *Sp^WAK2^-mCherry-HDEL*. Interestingly, *ProBnWRL1* preferentially drove gene expression in H-position cells, whereas *ProBnWRL2* preferentially drove gene expression in N-position cells (**Fig. S5c-d**). These results suggest that positional signals in the Arabidopsis root could be transmitted to cis-regulatory elements of *BnWRLs*.

We next investigated whether the identified *WRL*s have conserved functions. As expected, in Arabidopsis *wer-1* mutant plants, *ProAtWER:AtWER-GFP* construct was able to rescue the excessive root hair phenotype (ectopic N-position root-hair cells) and restore appropriate N-position cell expression of the target gene *AtGL2* (as shown by the *ProAtGL2:GUS* reporter) (**Figs. 4b-d,g, S14a-c; Dataset 1**). Further, GFP fluorescence signals suggested that the AtWER- GFP fusion proteins preferentially accumulated in N-position cells (**Fig. 4h-i**). We discovered that all tested *ProAtWER:RrWRL-GFP* and *ProAtWER:BnWRL-GFP* constructs also possessed the ability to restore expression of *AtGL2* (**Fig. S14c-h**) and rescue the mutant phenotype of *wer- 1* roots (**Figs. 4e-g, S15a-d; Dataset 1**), and the WRL-GFP fusion proteins also accumulated in N-position cells (**Figs. 4j- m,S15e-j**). These results suggested that the identified WRLs have conserved WER-like protein function in regulating expression of *AtGL2* and non-hair cell fate specification contributing to the Type III root epidermal cell patterns.

Interestingly, we noticed ectopic non-hair cell development and *ProAtGL2:GUS* activation in H-position in lines expressing *AtWER-GFP*, *RrWRL2-GFP*, and *BnWRL-GFP*s (**Fig. S14c,e,g,h**), accumulation of the WRL-GFP fusion proteins in H-position cells in lines expressing *BnWRL-GFP*s (**Figs. 4i-m,S15i-j**), and weaker rescue of the ectopic root hairs in N-position cells in lines expressing *BnWRL2-GFP* (**Fig. S15d, Dataset 1**). These variations suggest that these *WRL*s may be differently regulated when expressed in the Arabidopsis roots.

The absence of *WRL*s from the Type I species *C.sativus* led us to examine other Type I superrosid species to investigate whether there is an association between lack of this trait- specific MYB gene and the Type I pattern. *Pisum sativum* (Fabales) and *Linum usitatissimum* (Malpighiales), which adopt the Type I pattern (Clowes, 2000; Pemberton *et al*., 2001), were selected for analysis and we identified no WRL from these two species (**Fig. S16**). This result suggests that loss of the trait-specific MYB may account for the Type I pattern in the examined superrosid species.

### Identification and functional analysis of *CPC* homologs

In Arabidopsis roots, lateral inhibition mediated by CPC- related R3 MYB proteins plays an essential role in establishing the Type III pattern (Lee & Schiefelbein, 2002).

<u>CP</u>C-like (CPL) MYB proteins were identified from the selected species and the ML tree revealed a well-supported clade containing 7 *A.thaliana* proteins, 2 *R.crenulata* proteins, 2 *R.rosea* proteins, 2 *B.nivea* proteins, 1 *C.sativus* protein, and 1 *T*. *aralioides* protein (**Fig. 5a**) . *AtCPC*, *AtTRY*, *AtCPL3*, *AtETC1*, *RcCPL1*, *BnCPL1*, *BnCPL2*, *CsCPC*, and *TaCPL1* genes displayed shared synteny (**Fig. S3d**). We therefore refer to these *CPL*s as *AtCPC* orthologs. Interestingly, additional CPLs were identified from *R.crenulata*, *R.rosea*, *B.nivea*, and *T*. *aralioides*, which clustered in a clade sister to the AtCPC ortholog clade (**Fig. S17a**). Genes encoding these proteins were included in a synteny network without any *AtCPC* ortholog (**Fig. S3e**). We thus designated these genes as *AtCPC* paralogs. Sequence alignment of CPLs showed that they share a conserved R3 MYB domain, whereas AtCPC orthologs have Lysine (K)- and Arginine (R)-rich N and C termini not found in the AtCPC paralogs (**Fig. S18**).

**Fig. 5.**
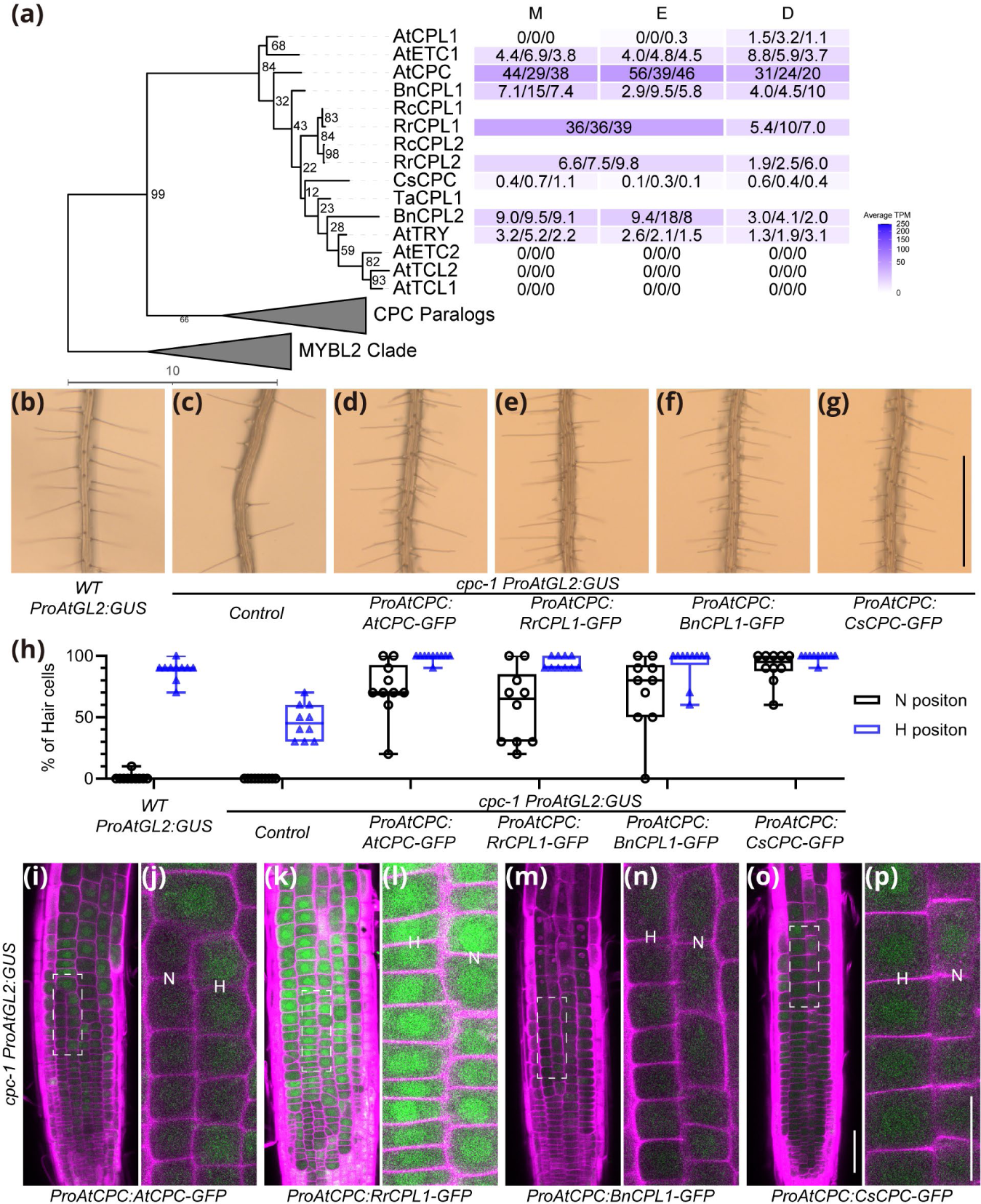
Identification and functional analysis of *CPL*s. (**a**) Left, maximum likelihood tree of CPL proteins. The tree is rooted to MYBL2 clade. The scale bar refers to amino acid substitutions per site. The CPC paralog clade is collapsed and is shown in **Fig. S17**. Right, gene expression levels in root developmental zones. TPM values of 3 RNA-seq replicates are shown in each box, and the average values are visualized by heatmaps after square root transformation. (**b**-**g**) Root hair phenotypes of Arabidopsis WT, *cpc-1* and *cpc-1* mutant bearing *ProAtCPC:CPL- GFP* constructs. Each line also carries *a ProAtGL2:GUS* transgene. All transgenic lines shown are representative T3 individuals. Bar=1mm. (**h**) Quantification of root epidermal cell phenotypes of the lines shown in (**b**-**g**). Root hair counting raw data and information of other independent lines are listed in **Dataset 1**. (**i**, **k**, **m**, **o**) Confocal images of roots from the lines shown in (**d**-**g**). Bar=50um. (**j**, **l**, **n**, **p**) Zoom in view of the regions highlighted by dash lines in (**i**, **k**, **m**, **o**). N-position cell files are labelled by N, and H-position cell files are labelled by H. Bar=25um. GFP signals are pseudocolored as green, CalcoFluor White signals are pseudocolored as magenta.

In Arabidopsis roots, *AtCPC, AtTRY and AtETC1* are expressed throughout all 3 developmental zones of the root, with *AtCPC* exhibiting the greatest transcript accumulation (**Fig. 5a; Dataset 4**). The *RrCPL1*, *RrCPL2, BnCPL1 and BnCPL2* transcripts were detected throughout all developmental zones of the roots with *RrCPL1* transcripts preferentially accumulating in the meristem/elongation zone (**Fig. 5a**; **Dataset 4, RrMEvsD**), whereas *CsCPC* transcripts were barely detected in *C.sativus* roots (**Fig. 5a**). The transcripts from *AtCPC* paralogs *BnCPL3*, *BnCPL4*, *BnCPL5*, and *RrCPL3* were also detected in the roots (**Fig. S17a**).

In the Arabidopsis root epidermis, *AtCPC* is transcribed and translated in N-position cells, and movement of AtCPC proteins from N-position cells to neighboring H-position cells leads to lateral inhibition (Lee & Schiefelbein, 2002; Wada *et al*., 2002; Koshino-Kimura *et al*., 2005; Kurata *et al*., 2005). We sought to investigate the regulation of *AtCPC* orthologs in the root epidermis. As expected, *ProAtCPC* was able to drive expression of the marker gene in N-position cells in WT Arabidopsis plants (**Fig. S5e**). We found that *ProBnCPL1* could similarly drive reporter gene expression in N-position cells in Arabidopsis roots (**Fig. S5f**), suggesting conservation of the cis-regulatory elements.

We next investigated functions of the identified *CPL*s by molecular complementation. As expected, a control *ProAtCPC:AtCPC-GFP* construct was able to restore root hair development in H-position cells in Arabidopsis *cpc-1* mutant plants. We also observed some ectopic root hair formation in N-position cells (**Fig. 5b-d,h; Dataset 1**), consistent with a previous study (Kurata *et al*., 2005). The GFP fluorescence signals showed that AtCPC-GFP proteins accumulate in nuclei of both N- and H-position cells (**Fig. 5i-j**), confirming mobility of AtCPC-GFP. In *cpc-1* mutant roots, *AtGL2* was ectopically expressed in H-position cells as shown by the *ProAtGL2:GUS* reporter (**Fig. S19a,b**). Unexpectedly, the *ProAtCPC:AtCPC-GFP* construct did not rescue the ectopic expression of *AtGL2* (**Fig. S19c**). Like *ProAtCPC:AtCPC- GFP*, we discovered that the *ProAtCPC:RrCPL1-GFP*, *ProAtCPC:BnCPL1-GFP*, and *ProAtCPC:BnCPL2-GFP* constructs also over-rescued the *cpc-1* mutant phenotype (**Figs. 5e,f,h, S17b,d; Dataset 1**). The CPL-GFP fusion proteins accumulated in both N- and H-position cells but with varied fluorescent signal (**Figs. 5k-p, S17e,f**), and the *ProAtGL2:GUS* reporter was activated in both N- and H- position cells (**Fig. S19d-f**). We also tested the one *CPL* identified from *C.sativus* (*CsCPC*) and made similar observations (**Figs. 5 g,h,o,p, S19h**). Together, these results suggest conserved functions of AtCPC ortholog proteins from these species.

We next analyzed members of the *AtCPC* paralog clade (**Fig. S17**). In molecular complementation experiments, the *ProAtCPC:BnCPL4-GFP* transgenic line exhibited restoration of root hair formation in H-position cells to varied extents (**Figs. S17c,d,S19g**; **Dataset 1**). The BnCPL4-GFP fluorescent signal was relative weak, but could be observed in nuclei of all root epidermal cells (**Fig. S17g,h**), suggesting that this protein could also move from N-position cells to H-position cells.

In Arabidopsis, *AtCPC* works redundantly with other orthologs in restricting trichome formation (Schellmann *et al*., 2002; Kirik *et al*., 2004a; Kirik *et al*., 2004b; Wang *et al*., 2008). Lines bearing the *ProAtCPC:AtCPC-GFP* transgenes lacked trichomes on their true leaves (**Fig. S20a-c**), consistent with a previous report (Kurata *et al*., 2005). Similarly, leaves of lines expressing *AtCPC* orthologs or paralog were also trichomeless (**Fig. S20d-h**). These results support the conclusion that the identified CPL proteins are functionally conserved across Type III species.

## Discussion

In this study, we identified and analyzed multiple superrosid species for the molecular basis of their root epidermal cell pattern. We discovered conservation in the GRN controlling the Arabidopsis position-dependent Type III pattern across multiple species from different lineages. Our results suggest that this is an ancient pattern-forming GRN that likely existed in the common ancestor of these diverse Type III species. Further, our analysis of Type I species suggests that their pattern resulted from loss or silencing of components of the conserved Type III GRN. Together, our work sheds light on the evolution of cell type patterns in angiosperms.

### Functional conservation and divergence of the GRN regulating root epidermal cell patterning

#### *RHD6* homologs

Our analyses of *RrRLB*s, *BnRLB*, and *CsRLB* suggest a conserved role for *RLB*s in promoting root hair initiation in both Type I and Type III species. This conclusion is consistent with prior studies of liverwort and moss *RHD6* orthologs, which suggested functional conservation of genes in this family across all land plants (Menand *et al*., 2007; Proust *et al*., 2016). Given the critical role of *RLB*s in triggering root hair formation, it may be that *RLB* gene expression represents the prime target of molecular mechanisms that regulate root epidermal cell type patterns.

It is notable that, in our cross-species complementation experiments, GFP-RLBs accumulated in nuclei of both N- and H-position cells when expressed by *ProAtRHD6* (**Fig. 2h-m**), but only H-position cells produced root hairs (**Fig. 2g; Dataset 1**). These observations are consistent with previous studies in which *RLB*s were ectopically expressed in the Arabidopsis N-position cells but did not induce ectopic root hair development (Lin *et al*., 2015; Kim & Dolan, 2016).

Interestingly, the GFP-RLBs seem to localize differently within nuclei of N- and H-position cells (**Fig. 2h-m**). These results suggest that functionality of RLBs in the root epidermis of Type III species may be dependent on factors that regulate their localization within nuclei.

#### *GL2* homologs

Our results demonstrate that GLHs from Type III species possess conserved functions when expressed in Arabidopsis (**Figs. 3,S7,S8)**, and our analysis of *ProBnGLH* (Figure S5a) indicates that *BnGLH* can respond to position-dependent factors like *AtGL2*. These findings suggest that *GLH*s play a conserved role in Type III species to direct non-hair cell differentiation in N-position cells.

It is notable that *CsGLH*s are expressed at very low levels in *C.sativus* roots, possibly explaining the lack of non-hair cells in this Type I species (**Fig. 3a; Dataset 4**).

Interestingly, our molecular complementation experiments showed that *CsGLH1* can rescue the Arabidopsis *gl2-1* mutant phenotypes in a dosage-dependent manner (**Figs. 2g,h,S6,S7F,S8F; Dataset 1**). This contrasts with a previous study in which *CsGLH1* (*CsGL2-LIKE*) driven by *ProCaMV35S* failed to rescue phenotypes of the Arabidopsis *gl2-8* mutant (Cai *et al*., 2020). This may be explained by the difference in promoters used, implying that rescue by *CsGLH1* may require a particular level and/or developmental timing of gene expression.

#### *TTG1* homologs

We identified *TLW*s from both Type I and Type III species which have shared synteny and encode proteins with high amino acid similarity (**Fig. S9a,c,S10**).

The *TLWs* from Type III species were expressed in roots in a similar pattern, whereas *CsTLW* was expressed at a relatively low level (**Supplemental Fig. 9A**; Dataset 4). This suggests that different expression/accumulation levels of TLWs, rather than divergent protein functions, may be associated with the Type III versus Type I patterns.

#### *GL3* homologs

We identified GLB proteins from Type III species *R.rosea* and *B.nivea*, but not from Type I species *C.sativus* or the basal eudicot *T.aralioide* (**Fig.S 9b**). *GL3* homologs from multiple species have been shown to complement the Arabidopsis *gl3 egl3* mutant phenotypes (Zhang & Hülskamp, 2019). Considering the sequence conservation found in the identified full length GLBs (**Fig. S11**), we suggest that these have conserved protein functions.

Our phylogenetic analysis also revealed a well-supported clade sister to the GLB clade, which includes AtMYC1 homologs (MLBs) from all selected species (**Fig. S9b).**

Interestingly, it was reported that *AtGL3* and *AtEGL3* can complement Arabidopsis *myc1* mutant phenotypes, whereas *AtMYC1* cannot complement the *gl3 egl3* mutant phenotypes (Zhao *et al*., 2012). In this study, we demonstrated that *MLB*s and *GLB*s have shared synteny (**Fig. 9d**). Thus, *GLB*s may have arisen from duplication of ancient *MLB*s, followed by subfunctionalization of the genes in these two clades.

#### *WER* homologs

We identified WER homologs (WRLs) from the Type III species *R.rosea* and *B.nivea*, as well as the basal eudicot *T.aralioide*, but not from the Type I species *C.sativus*, *P.sativum* or *L.usitatissimum* (**Figs. 4a,S16**), and most of these identified *WRL* genes have shared synteny (**Fig. S3c**). Thus, the absence of *WRL* from Type I species can be explained by the loss of *WRL* genes during speciation. The correlation between the presence of a *WRL* gene and the Type III pattern in these plant species suggests a critical role for *WRL*s for this cell type pattern.

Our cross-species complementation experiments showed that all WRLs from Type III species we investigated have the fundamental ability to rescue the Arabidopsis *wer-1* mutant phenotype (**Figs. 4g,S15**) through activation of *AtGL2* transcription in N-position cells (**Fig. S14**). Interestingly, we also noticed that despite the same *ProAtWER* cis-regulatory sequence was used to drive expression of *WRL-GFP*s, the root hair phenotypes, accumulation patterns of WRL-GFP proteins, and activation of the *ProAtGL2:GUS* reporter varied to some extent among the transgenic lines (**Figs. 4g- m,S14d-j,S15; Dataset 1**). These results suggest that WRLs from different species may have somewhat different activity or stability in the Arabidopsis root epidermis.

#### *CPC* homologs

We identified *CPC* homologs (*CPL*s) from all species we examined and categorized these genes into *CPC* ortholog and *CPC* paralog groups (**Figs. 5a,S17a**).

Our root zonation RNA-seq results showed that *AtCPC* orthologs from the Type III species *R.rosea* and *B.nivea* were expressed in a pattern similar to *AtCPC*, whereas *CsCPC* was expressed at a low level (**Fig. 5a; Dataset 4**). Furthermore, cis-regulatory elements of *BnCPL1* were shown to respond to cell-position specific factors in the Arabidopsis root epidermis similar to *ProAtCPC* (**Fig. S5e,f**), and our molecular complementation experiments showed that all tested AtCPC ortholog proteins have conserved functions (**Fig. 5h; Dataset 1**). Together, these results suggest the ancestral gene of *AtCPC* orthologs was responsible for mediating lateral inhibition for the Type III root epidermal cell pattern. Presumably, *AtCPC* orthologs underwent duplication and divergence of cis-regulatory elements, resulting in a varied number of *CPL*s with different expression patterns.

We identified *AtCPC* paralogs from the Type III species *R.rosea* and *B.nivea*, and the basal eudicot *T.aralioide* (**Fig. S17a**). AtCPC paralog proteins have also recently been reported in *Camellia sinensis* (Wakamatsu *et al*., 2021), but overlooked in most similar studies, potentially due to limited number of species and proteins analyzed. Although we found that the *AtCPC* paralogs were expressed in roots (**Fig. S17a**), and could rescue the *cpc-1* mutant phenotype (**Fig. S17c**), they show evidence of divergence from the *AtCPC* orthologs, in their expression profiles (**Fig. S17a)** and phenotypes of the *ProAtCPC:BnCPL4-GFP* line (**Fig. S17c,d; Dataset 1**). The lack of the short segments rich in K and R residues found in the N- and C- termini of AtCPC proteins (**Fig. S18**) may account for the diverged functions of the AtCPC paralog proteins.

### Molecular evolution of root epidermal cell patterns in superrosids

The discovery of the Type III pattern in multiple scattered species across the superrosids clade (Clowes, 2000; Pemberton *et al*., 2001), and our molecular analysis of patterning gene homologs reported here allow us to reconsider evolution of the Type III pattern at the scale of the whole superrosids clade.

We propose the following model. Considering the conserved function of *RLB*s in regulating tip growth across land plants (Menand *et al*., 2007; Proust *et al*., 2016), we propose in an ancestral euphyllophytes plant, *RLB*s were activated in all root epidermal cells and were sufficient to initiate root hair development, resulting in a Type I pattern in which all epidermal cells differentiate into root hair cells (**Fig. 6a**).

**Fig. 6.**
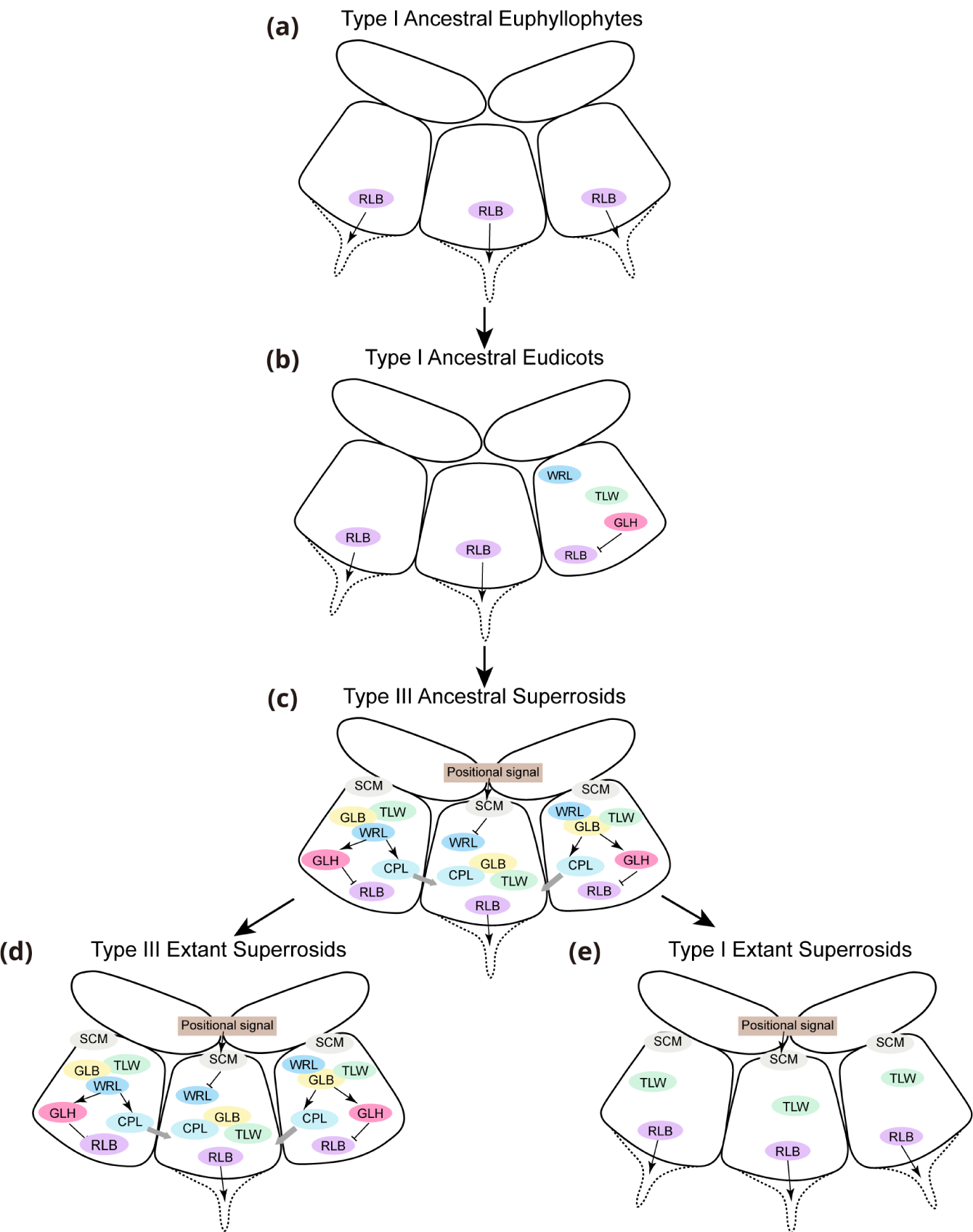
Model of evolution of root epidermal cell patterns in Superrosids

We next propose that a Type I ancestral eudicot evolved that was able to generate some non-hair epidermal cells via the production of GLH proteins in some (random) root epidermal cells, which inhibited the expression of the *RLB* genes in those cells (**Fig. 6b**).

Next, the ancestral eudicot species underwent molecular changes that converted the random activation of *GLH* expression to position-dependent activation via the preferential production of an MBW complex in the N-position cells. Multiple events would likely have occurred to generate this ancestral Type III species (**Fig. 6c**), including the regulation of *WRL* expression by a positional signaling pathway mediated by the SCM receptor. The WRL proteins also gained the ability to bind to specific cis-regulatory elements of *CPLs* and *GLHs*, allowing the MBW complex to activate these genes in N-position cells. Once this mechanism was established, it was inherited by extant Type III superrosid species like *R.rosea*, *B.nivea*, and *A.thaliana* (**Fig. 6d**).

Our analysis of the Type I species *C.sativus*, *P.sativum*, *L.usitatissimum* and genetic studies in *A.thaliana* (Galway *et al*., 1994; Masucci *et al*., 1996; Lee & Schiefelbein, 1999; Bernhardt *et al*., 2003) suggest that mutations in conserved patterning genes result in loss of the Type III pattern. Thus, we propose that the many extant lineages of Type I species across the superrosids group may have arisen in a similar manner via loss or silencing of *WRLs* and/or other patterning genes (**Fig. 6e**).

## Acknowledgements

We thank Erik Nielsen, Cora MacAlister, Regina Baucom, Jianming Li, Kook Hui Ryu, Wenjia Wang, and Fei Zhou (University of Michigan, Ann Arbor) for suggestions and helpful discussions. We thank Gregg Soboccinski and Leibin Wang (University of Michigan, Ann Arbor) for helpful discussions about microscopy. We thank Michael Palmer (The University of Michigan Matthaei Botanical Gardens, Ann Arbor) for helpful discussions about plant growth and pest management. This work was supported by National Science Foundation grants IOS-1923589 and IOS-2127485 and Department of Energy grant DE-SC0020358.

## Author contributions

Y.Z. and J.S. designed the research, performed the experiments, analyzed the data, and wrote the article.

## Data availability

The RNA sequencing data have been deposited in National Center for Biotechnology Information Bioproject database under the accession number PRJNA913166.

## Supporting Information

Fig. S1 *R.rosea* and *B.nivea* root images.

Fig. S2 Sequence alignment of RLBs.

Fig. S3 Microsynteny networks of *RLB*s, *GLH*s, *WRL*s, and *CPL*s.

Fig. S4 Sequence alignment of GLHs.

Fig. S5 Confocal images of Arabidopsis WT roots expressing *SpWAK2-mCherry-HDEL* controlled by different cis-regulatory elements.

Fig. S6 Phenotypes of a transgenic line with multiple *ProAtGL2:GFP-CsGL2* insertions.

Fig. S7 Trichome phenotypes of lines bearing *ProAtGL2:GFP-GLH* constructs.

Fig. S8 Seed mucilage phenotypes of lines bearing *ProAtGL2:GFP-GLH* constructs.

Fig. S9 Identification of *TTG1* and *GL3*/*EGL3* homologs.

Fig. S10 Sequence alignment of TLWs.

Fig. S11 Sequence alignment of full length GLBs, MLBs, and MGBs.

Fig. S12 Sequence alignment of truncated GLBs and MLBs.

Fig. S13 Sequence alignment of WRLs.

Fig. S14 Expression of *ProAtGL2:GUS* transcriptional reporter in Arabidopsis lines bearing *ProAtWER:WRL-GFP* constructs.

Fig. S15 Functional analysis of additional *WRL*s.

Fig. S16 Absence of *WRL*s from Type I species *L.usitatissimum* and *P.sativum*.

Fig. S17 Identification and functional analysis of additional *CPL*s.

Fig. S18 Sequence alignment of CPLs.

Fig. S19 Expression of *ProAtGL2:GUS* transcriptional reporter in Arabidopsis lines bearing *ProAtCPC:CPL-GFP* constructs.

Fig. S20 Trichome phenotypes of lines bearing *ProAtCPC:CPL-GFP* constructs.

**Methods S1** Root hair counting assay.

**Methods S2** Plastic embedding and transverse sectioning of plant roots.

**Methods S3** Generation of plasmids.

**Methods S4** RNA Isolation and sequencing.

**Methods S5** RNA-seq analysis.

**Methods S6** Phylogenetic analysis.

**Dataset 1** Raw data from root hair counting assay.

**Dataset 2** Primers and nucleotides used in this study.

**Dataset 3** Information about the genome and transcriptome resources used in this study and gene accessions.

**Dataset 4** Root transcript profiles of *A.thaliana*, *B.nivea*, *R.rosea*, and *C.sativus*.

