## Supplementary Figures and Methods for "A conserved gene regulatory network controls root epidermal cell patterning in superrosid species"

Yan Zhu and John Schiefelbein*
Department of Molecular, Cellular, and Developmental Biology, University of Michigan, Ann Arbor, Michigan 48109

* Author for correspondence:

John Schiefelbein

ORCIDs:

Yan Zhu: https://orcid.org/0000-0002-2655-6766

John Schiefelbein: https://orcid.org/0000-0002-0560-5872

The following Supporting Information is available for this article:

**Fig. S1** *R.rosea* and *B.nivea* root images.

**Fig. S2** Sequence alignment of RLBs.

**Fig. S3** Microsynteny networks of *RLB*s, *GLH*s, *WRL*s, and *CPL*s.

**Fig. S4** Sequence alignment of GLHs.

**Fig. S5** Confocal images of Arabidopsis WT roots expressing *SpWAK2-mCherry-HDEL* controlled by different cis-regulatory elements.

**Fig. S6** Phenotypes of a transgenic line with multiple *ProAtGL2:GFP-CsGL2* insertions.

**Fig. S7** Trichome phenotypes of lines bearing *ProAtGL2:GFP-GLH* constructs.

**Fig. S8** Seed mucilage phenotypes of lines bearing *ProAtGL2:GFP-GLH* constructs.

**Fig. S9** Identification of *TTG1* and *GL3*/*EGL3* homologs.

**Fig. S10** Sequence alignment of TLWs.

**Fig. S11** Sequence alignment of full length GLBs, MLBs, and MGBs.

**Fig. S12** Sequence alignment of truncated GLBs and MLBs.

**Fig. S13** Sequence alignment of WRLs.

**Fig. S14** Expression of *ProAtGL2:GUS* transcriptional reporter in Arabidopsis lines bearing *ProAtWER:WRL-GFP* constructs.

**Fig. S15** Functional analysis of additional *WRL*s.

**Fig. S16** Absence of *WRL*s from Type I species *L.usitatissimum* and *P.sativum*.

**Fig. S17** Identification and functional analysis of additional *CPL*s.

**Fig. S18** Sequence alignment of CPLs.

**Fig. S19** Expression of *ProAtGL2:GUS* transcriptional reporter in Arabidopsis lines bearing *ProAtCPC:CPL-GFP* constructs.

**Fig. S20** Trichome phenotypes of lines bearing *ProAtCPC:CPL-GFP* constructs.

**Methods S1** Root hair counting assay

**Methods S2** Plastic embedding and transverse sectioning of plant roots

**Methods S3** Generation of plasmids

**Methods S4** RNA isolation and sequencing

**Methods S5** RNA-seq analysis

**Methods S6** Phylogenetic analysis

**Dataset 1** Raw data from root hair counting assay.

**Dataset 2** Primers and nucleotides used in this study.

**Dataset 3** Information about the genome and transcriptome resources used in this study and gene accessions.

**Dataset 4** Root transcript profiles of *A.thaliana*, *B.nivea*, *R.rosea*, and *C.sativus*.

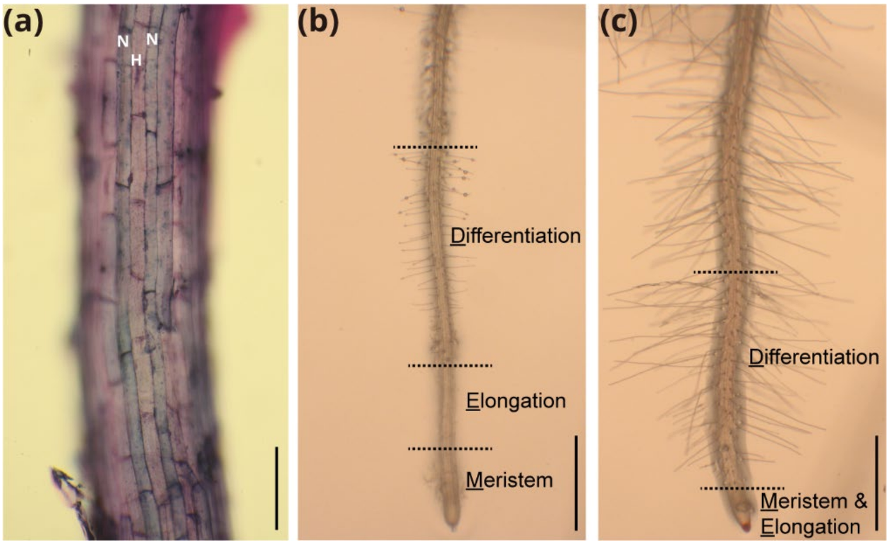


**Figure S1**
*R.rosea* and *B.nivea* root images. (**a**) Representative *B.nivea* primary root elongation zone stained with toludine blue. N-position cell files are labeled by N, and H-position cell file is labelled by H. Bar=100um. (**b** and **c**) Representative primary roots of *B.nivea* and *R.rosea*, respectively. The developmental zones collected for RNA sequencing are separated by the dash lines. Meristem and elongation zones were defined as hairless root tips. For *B.nivea*, the lower 40%-50% portion of the hairless root region was assigned as meristem zone, the remaining was assigned as elongation zone. For *R.rosea*, the meristem and elongation zones were combined to yield a sufficient amount of RNA. The differentiation zone was defined as the root segment from where root hairs start to emerge to the point where root hairs reach their maximum length. Bar=1mm

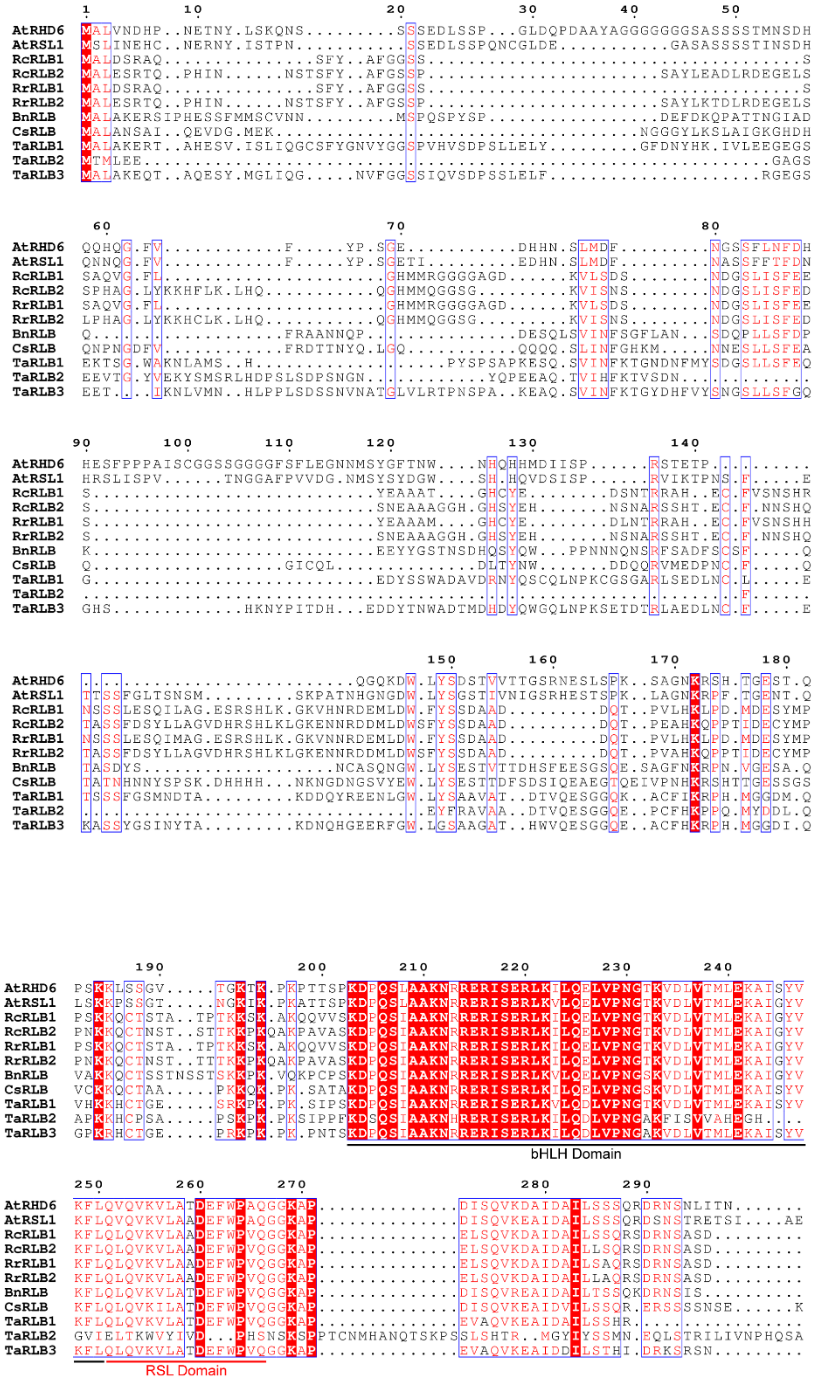

**Figure S2.**
Sequence alignment of RLBs. Conserved amino acids among some of the RLBs (>70%) are colored as red. Conserved amino acids among all RLBs are colored as white and shaded with red. Conserved regions shared by RLBs (>70%) are labelled by blue squares. The bHLH domain and RSL domain are highlighted.

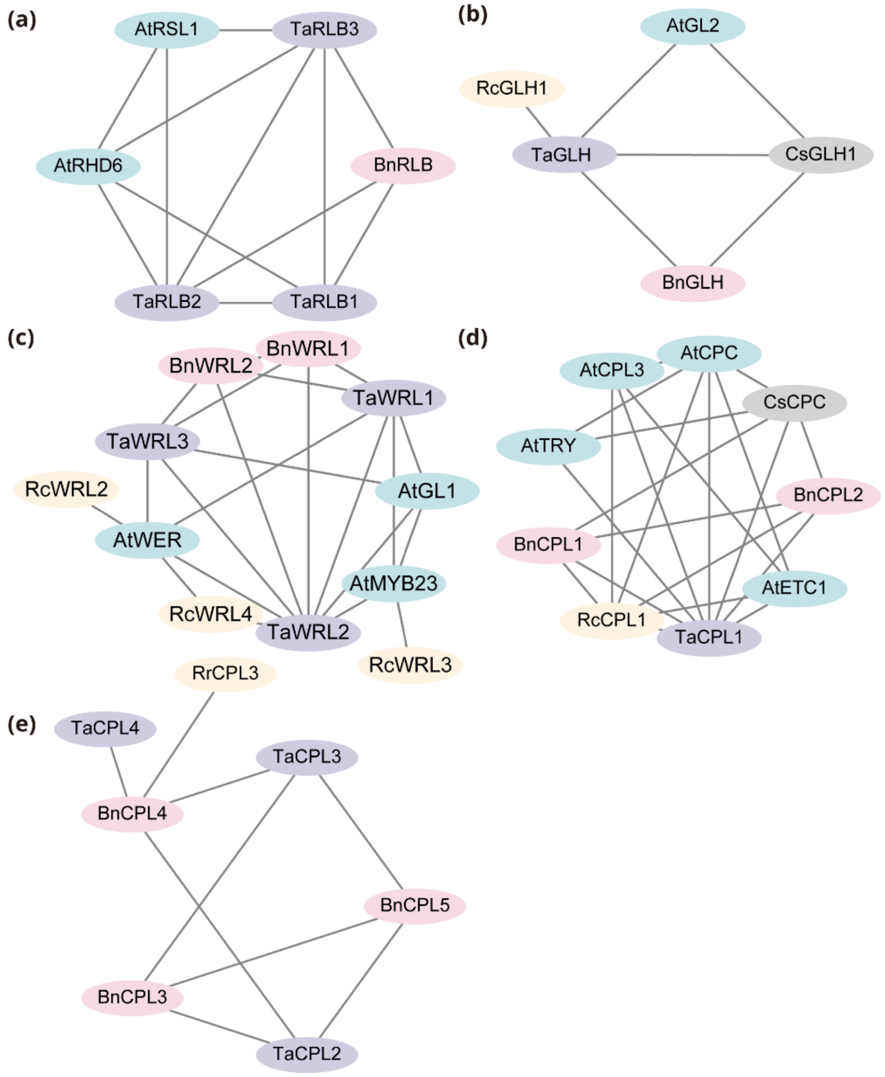


**Figure S3**
Microsynteny networks of *RLB*s, *GLH*s, *WRL*s, and *CPL*s. Nodes represent gene loci encoding corresponding proteins. Edges linking two nodes suggest syntenic connection of the two genes. Length of the edges are not informative.

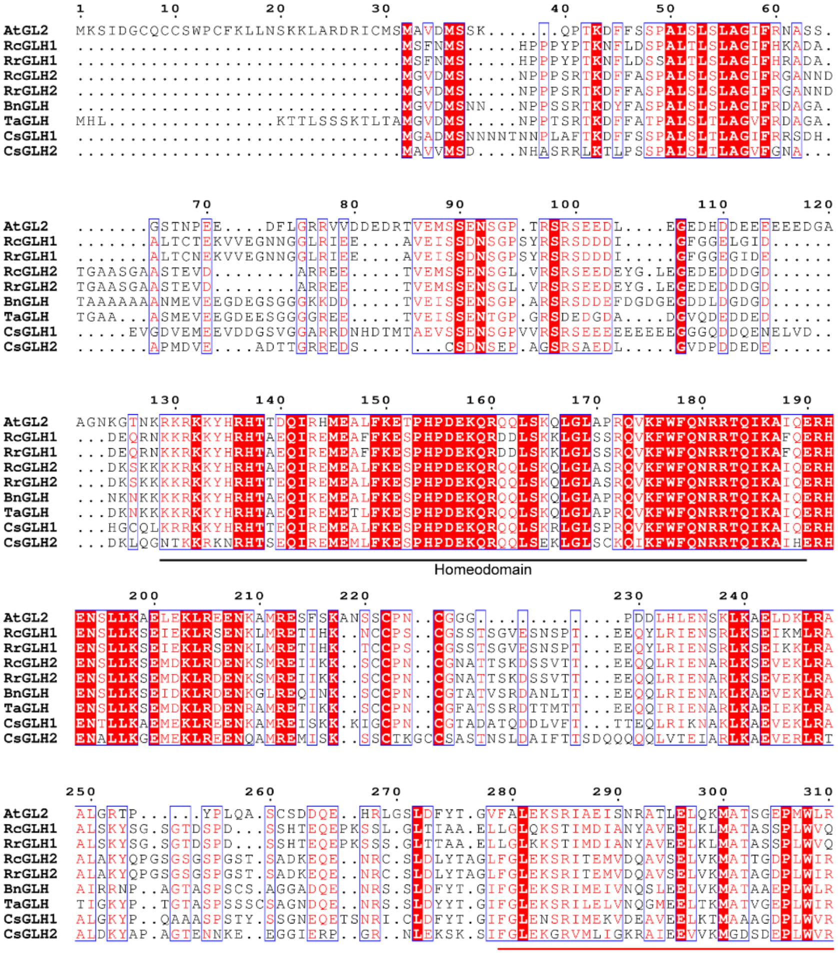
AtGL2
RcGLH1
RrGLH1
RcGLH2
RrGLH2
BnGLH
TaGLH
CsGLH1
CsGLH2
Homeodomain

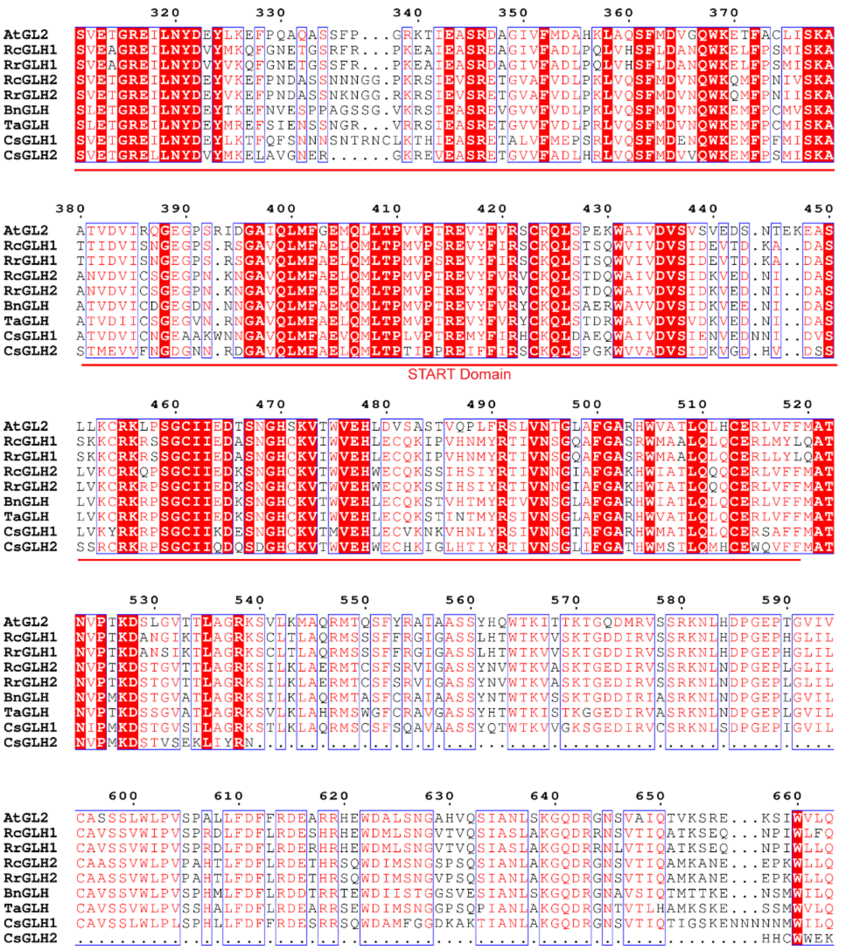
AtGL2
RcGLH1
RrGLH1
RcGLH2
RrGLH2
BnGLH
TaGLH
CsGLH1
CsGLH2
START Domain

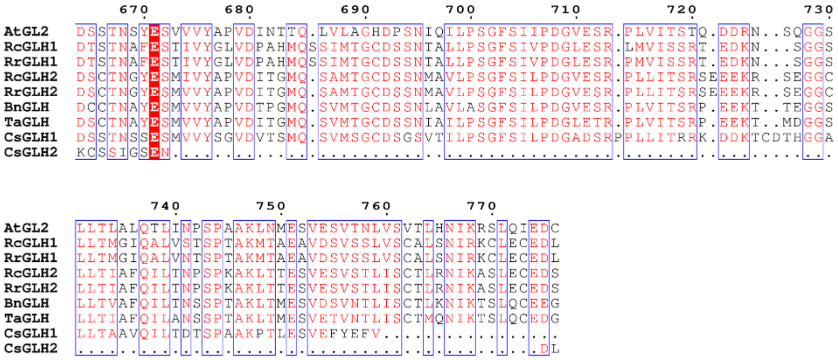


**Figure S4**

Sequence alignment of GLHs. Conserved amino acids among some of the GLHs (>70%) are colored as red. Conserved amino acids among all GLHs are colored as white and shaded with red. Conserved regions shared by GLHs (>70%) are labelled by blue squares. The Homeodomain and START domain are highlighted.

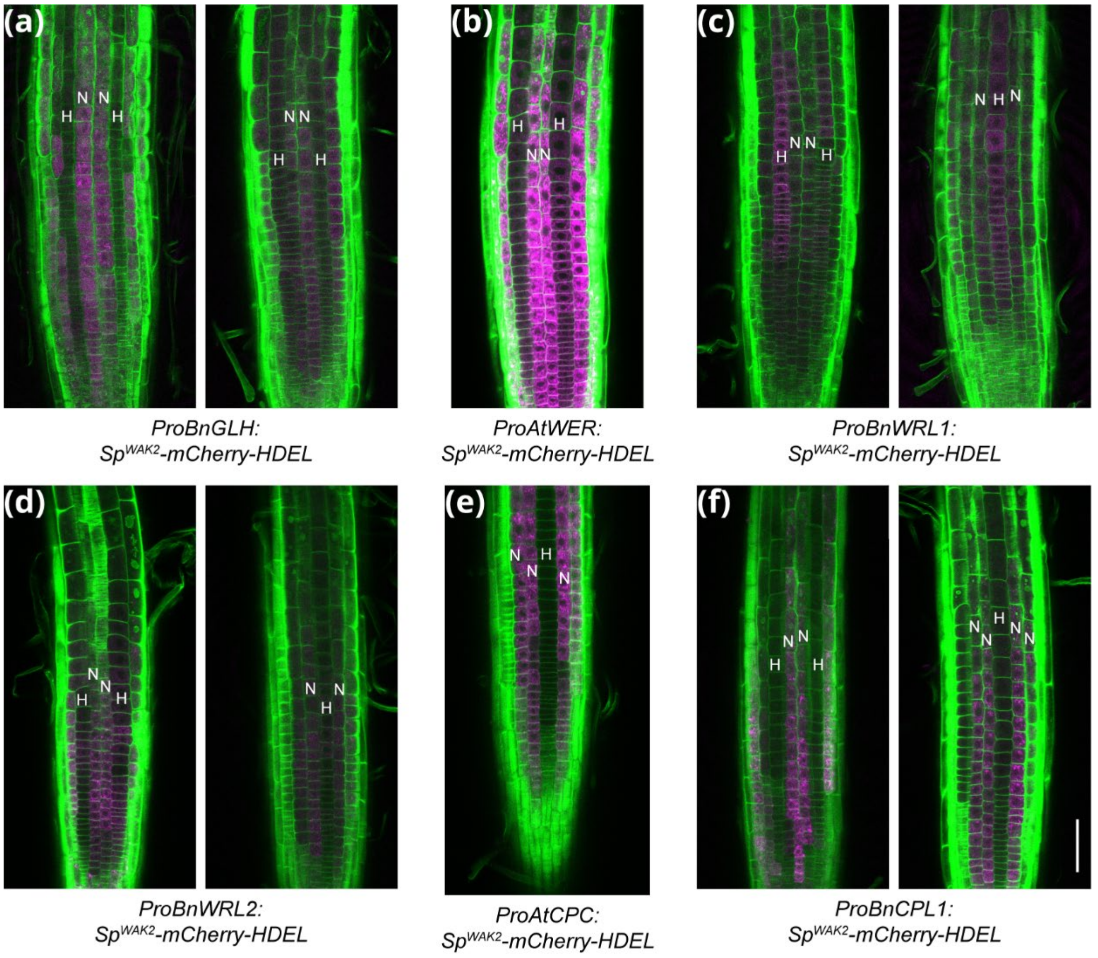


**Fig. S5**

Confocal images of Arabidopsis WT roots expressing *SpWAK2-mCherry-HDEL* controlled by different cis-regulatory elements. Seedling roots of independent homozygous T3 plants were imaged. N-position cell files are labelled with N, and H-position cell files are labelled with H. Intensity and brightness of (**a**) and (**c**) were adjusted differently to the other panels due to weak signals. mCherry signals are pseudocolored as magenta, Calcofluor White signals are pseudoclored as green. Bar=50um

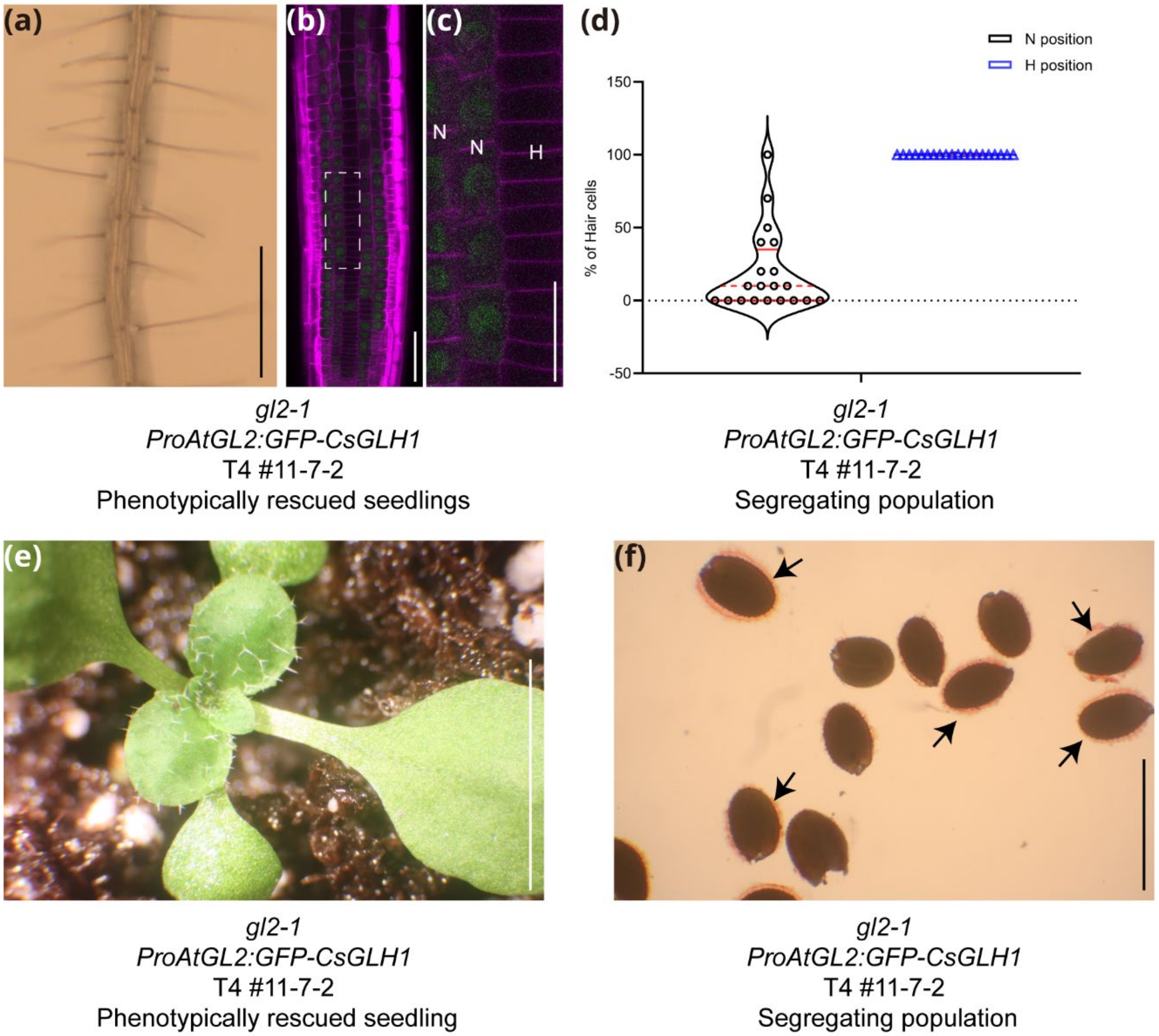


**Fig S6**

Phenotypes of a transgenic line with multiple *ProAtGL2:GFP-CsGL2* insertions. (**a**) Root hair phenotype of a T4 seedling showing rescued *gl2-1* mutant phenotype. Bar=1mm. (**b**) Confocal image of a T4 seedling showing rescued *gl2-1* mutant phenotype. Bar=50um. (**c**) Zoom in view of the region highlighted by dash lines in (**b**). N-position cell files are labelled by N, and H-position cell file is labelled by H. Bar=25um. GFP signals are pseudocolored as green, CalcoFluor White signals are pseudocolored as magenta. (**d**) Quantification of root epidermal cell phenotypes in a phenotypically segregating T4 population. The red line refers to median. The interquartile region is flanked by red dash lines. Root hair counting raw data is listed in **Dataset 1**. (**e**) Trichome phenotype of a T4 seedling showing rescued *gl2-1* phenotype. Bar=5mm. (**f**) Seed mucilage production in a phenotypically segregating T4 population. The arrows point to seeds producing mucilage. Bar=5mm.

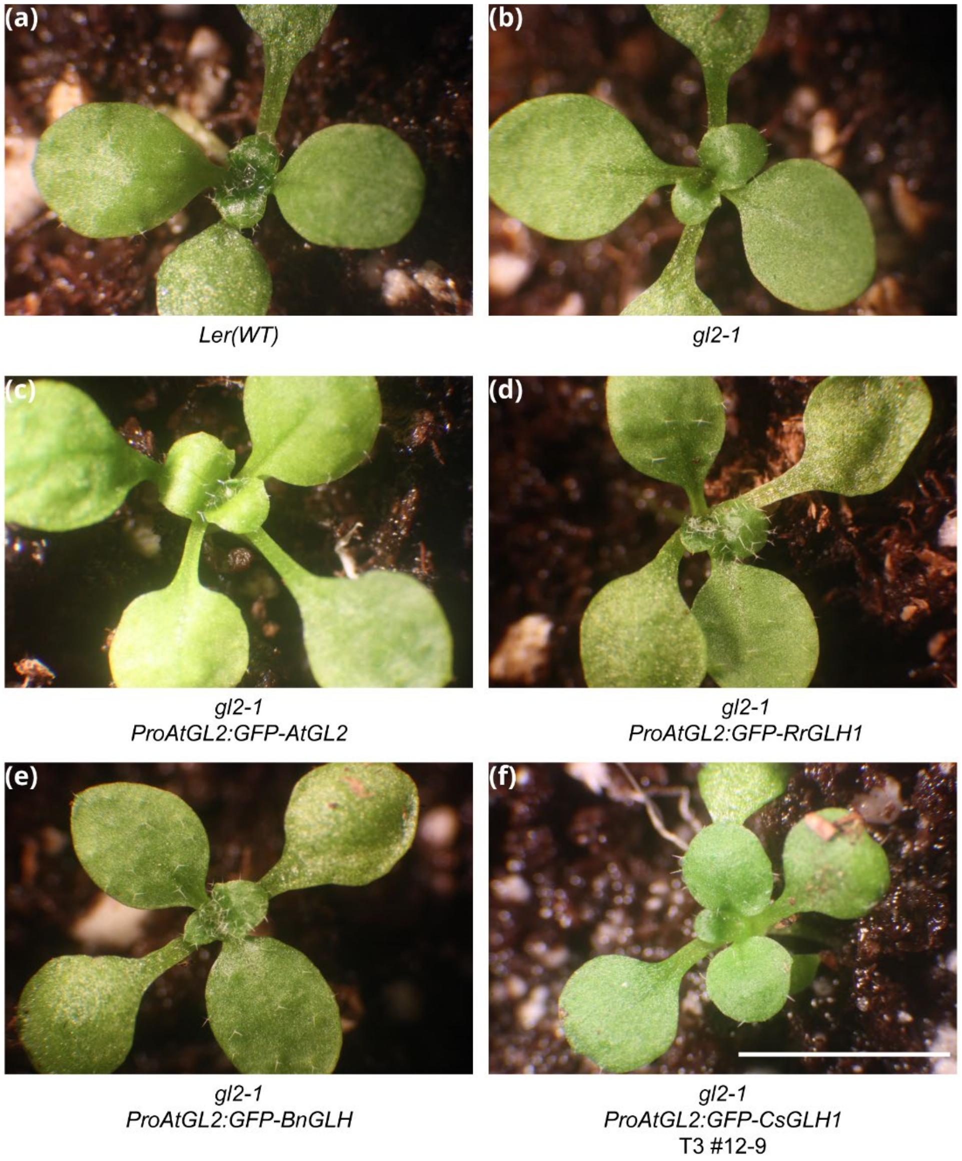


**Figure S7**

Trichome phenotypes of lines bearing *ProAtGL2:GFP-GLH* constructs. All transgenic lines shown are representative T3 individuals. Bar=5mm.

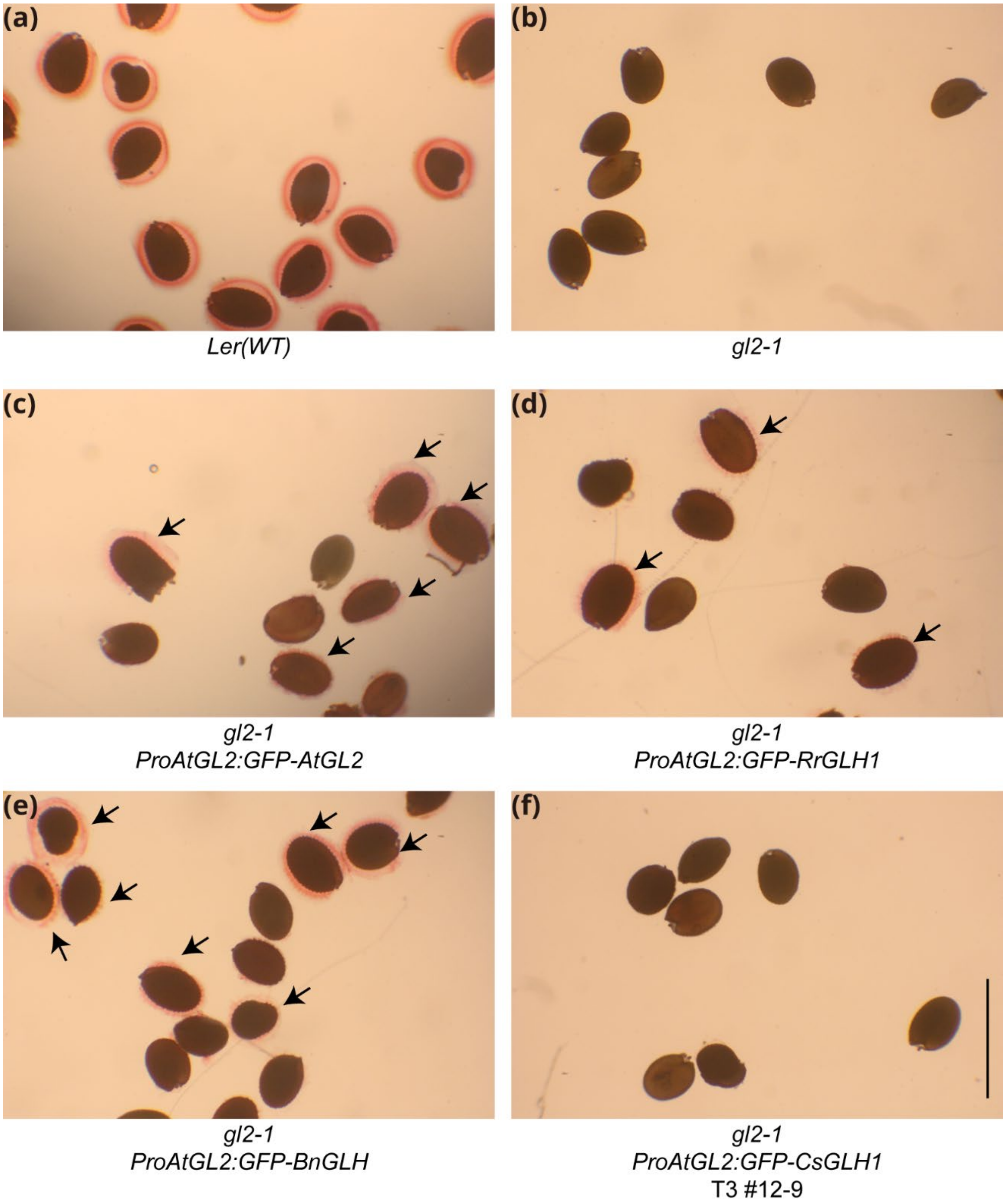


**Figure S8**
Seed mucilage phenotypes of lines bearing *ProAtGL2:GFP-GLH* constructs. In (**c**-**e**), the arrows point to seeds producing mucilage. Bar=5mm.

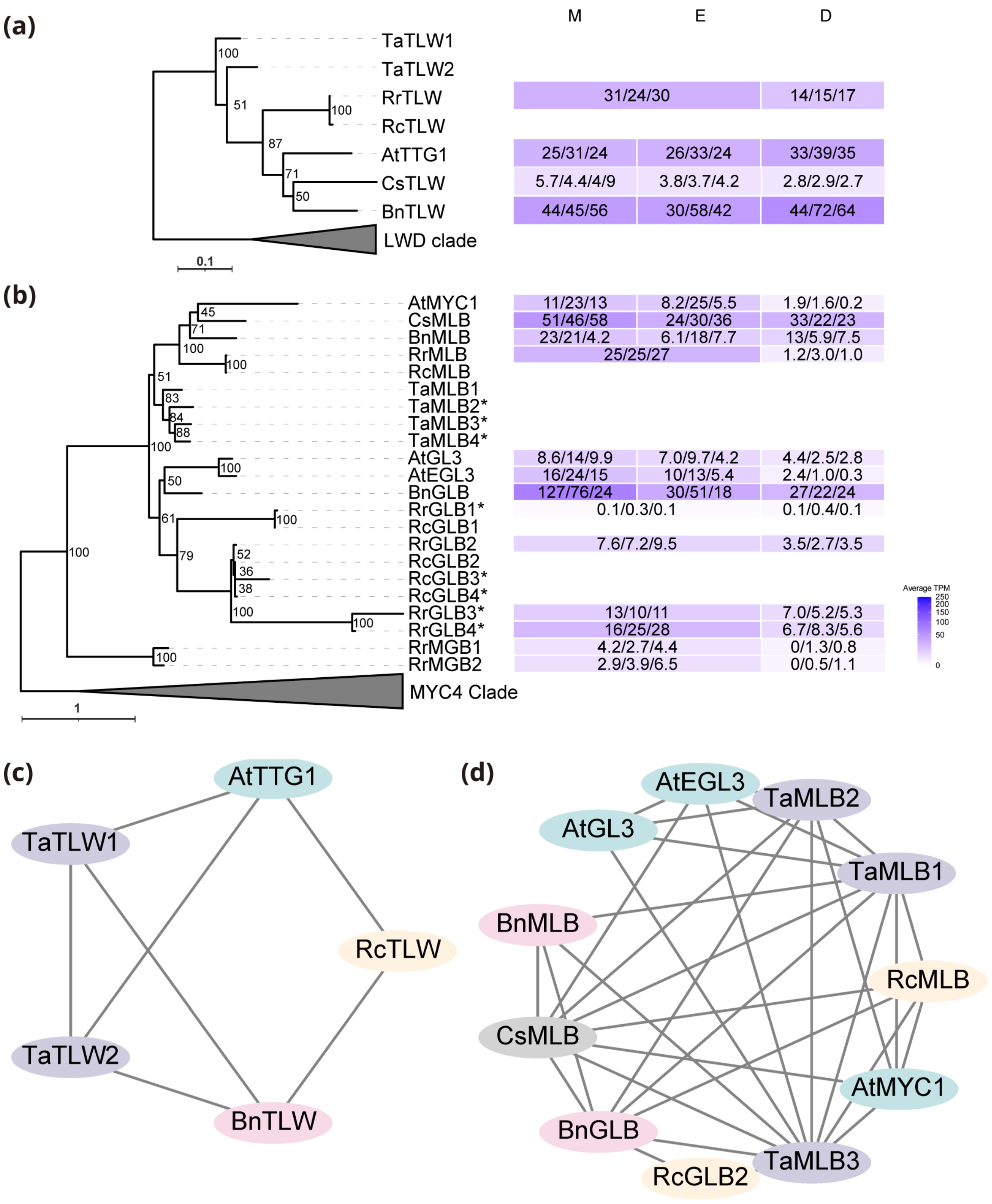


**Figure S9**

Identification of *TTG1* and *GL3*/*EGL3* homologs. (**a**) Left, maximum likelihood tree of TLWs. The tree was rooted to the LWD clade. The scale bar refers to amino acid substitutions per site. Right, gene expression levels in root developmental zones. TPM values of 3 RNA-seq replicates are shown in each box, and the average values are visualized by heatmaps after square root transformation. (**b**) Left, maximum likelihood tree of MLBs and GLBs. The tree was rooted to the MYC4 clade. The scale bar refers to amino acid substitutions per site. Right, gene expression levels in root developmental zones. TPM values of 3 RNA-seq replicates are shown in each box, and the average values are visualized by heatmaps after square root transformation. *, truncated proteins. (**c**) Microsynteny network of *TLW*s. (**d**) Microsynteny network of *MLB*s and *GLB*s.

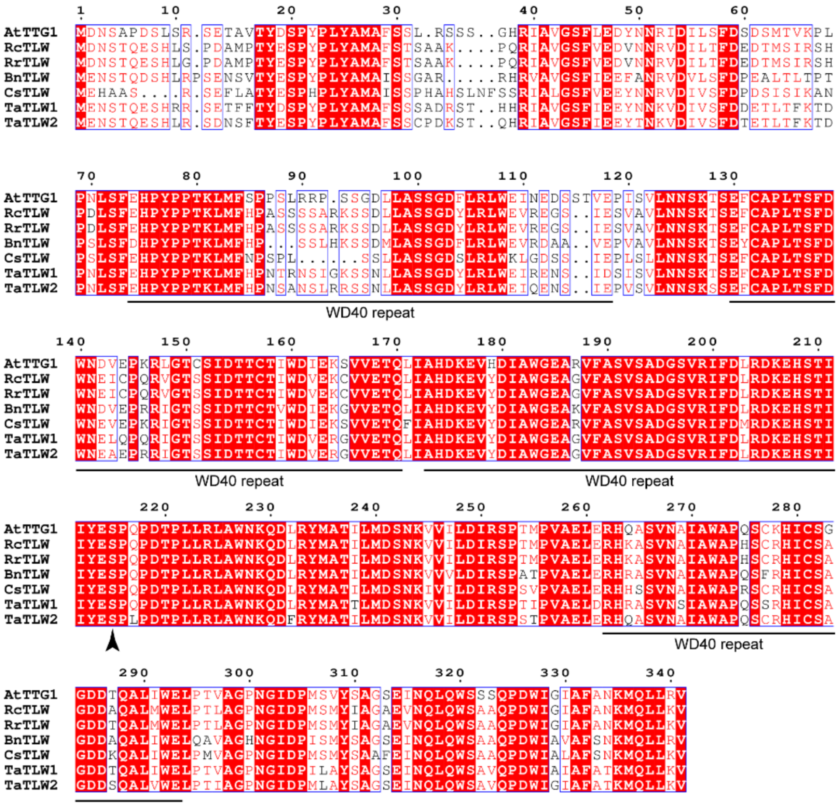


**Figure S10**

Sequence alignment of TLWs. Conserved amino acids among some of the TLWs (>70%) are colored as red. Conserved amino acids among all TLWs are colored as white and shaded with red. Conserved regions shared by TLWs (>70%) are labelled by blue squares. The WD repeats are highlighted. The arrowhead points to the amino acid that can be phosphorylated in AtTTG1.

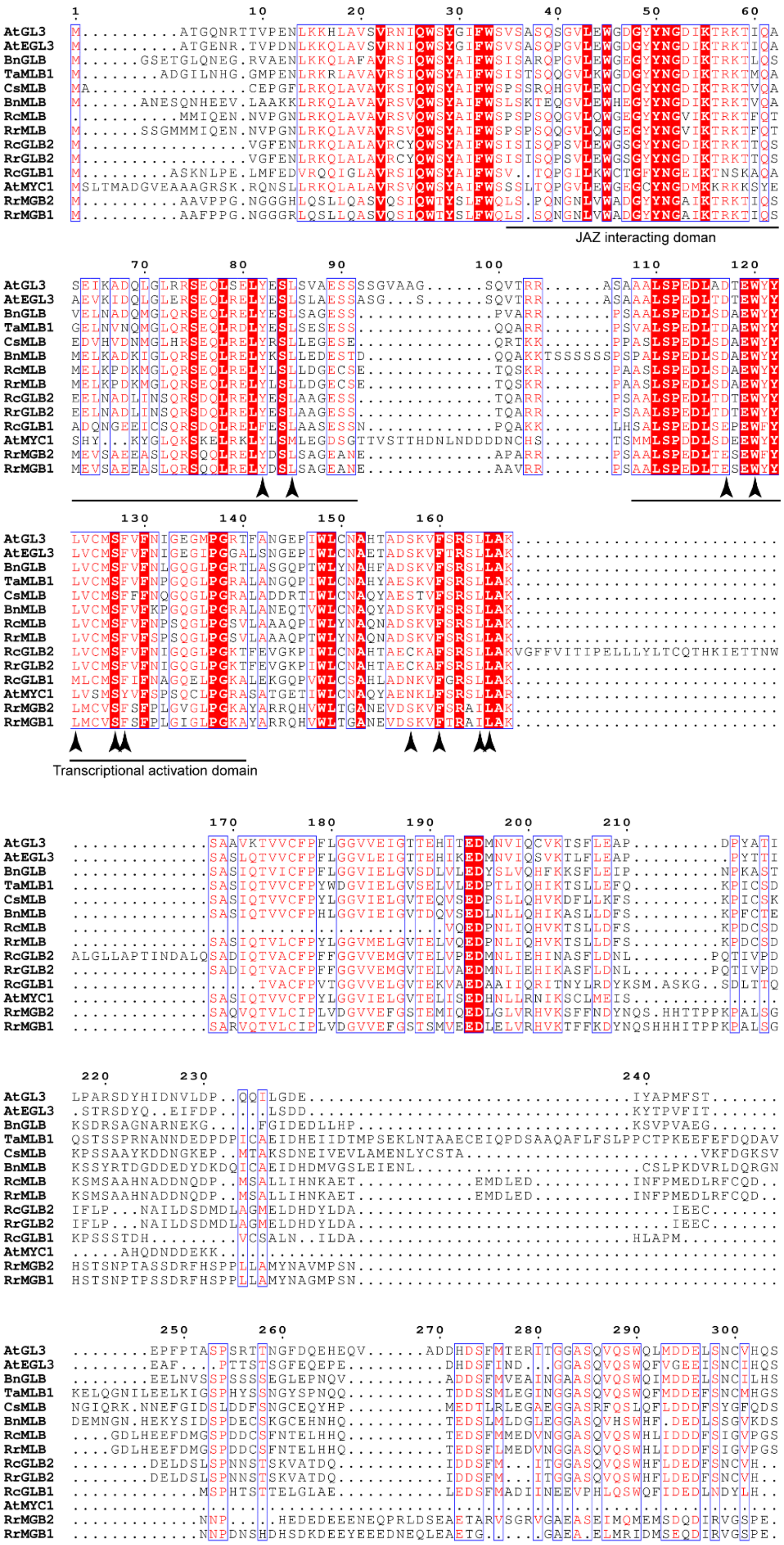
JAZ interacting doman
Transcriptional activation domain

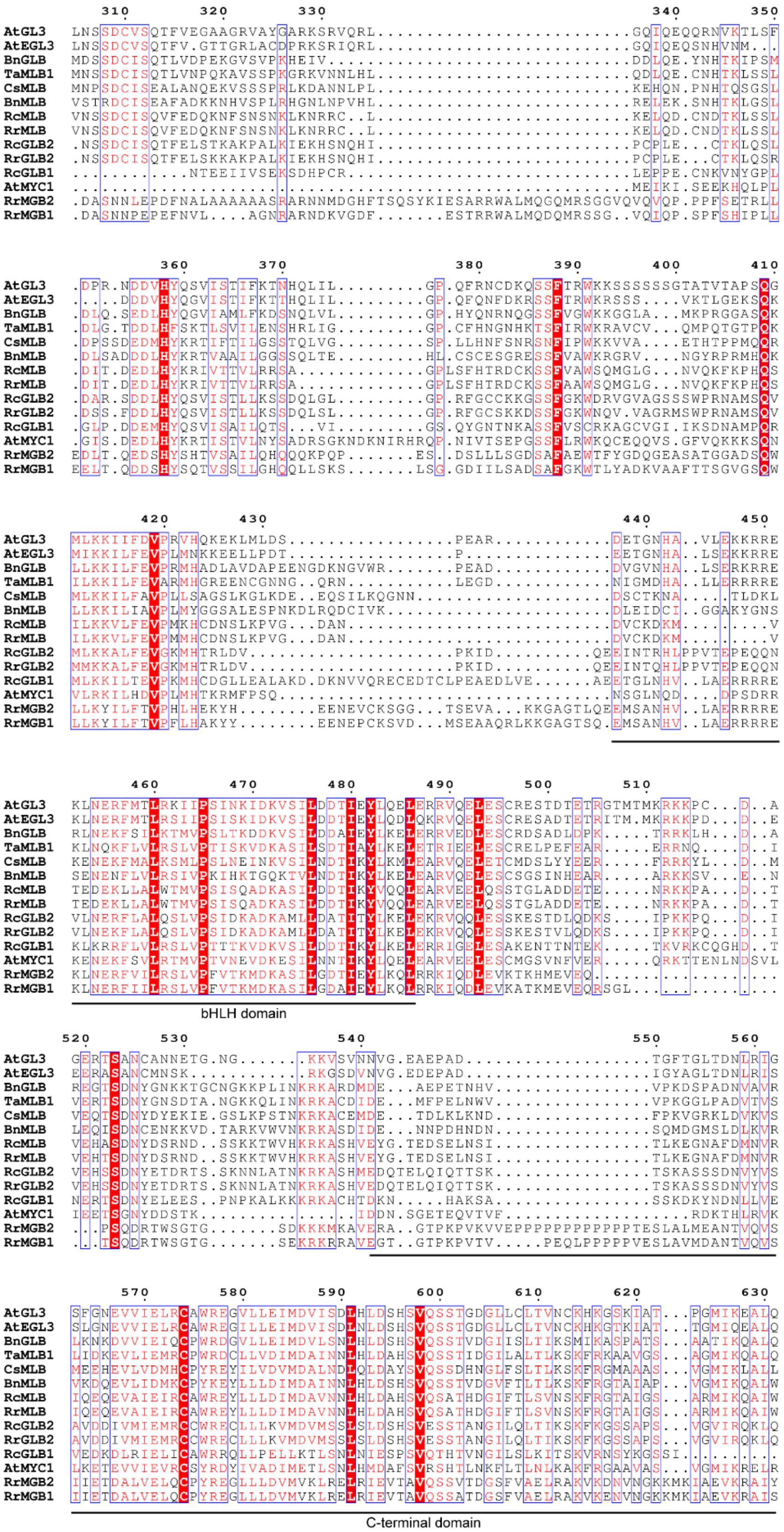
AtGL3
AtEGL3
BnGLB
TaMLB1
CsMLB
BnMLB
RcMLB
RrMLB
RcGLB2
RrGLB2
RcGLB1
AtMYC1
RrMGB2
RrMGB1
bHLH domain
C-terminal domain

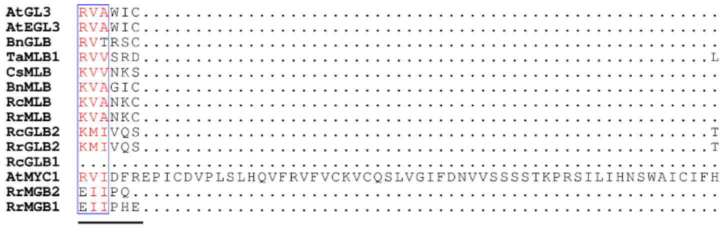


**Figure S11**

Sequence alignment of full length GLBs, MLBs, and MGBs. Conserved amino acids among some of the GLBs, MLBs, and MGBs (>70%) are colored as red. Conserved amino acids among all GLBs, MLBs, and MGBs are colored as white and shaded with red. Conserved regions shared by GLBs, MLBs, and MGBs (>70%) are labelled by blue squares. The JAZ interacting domain (JID), Transcriptional activation domain (TAD), bHLH domain, and C-terminal domain are highlighted. The arrowheads point to the residues mediating interaction with R2R3 MYB proteins.

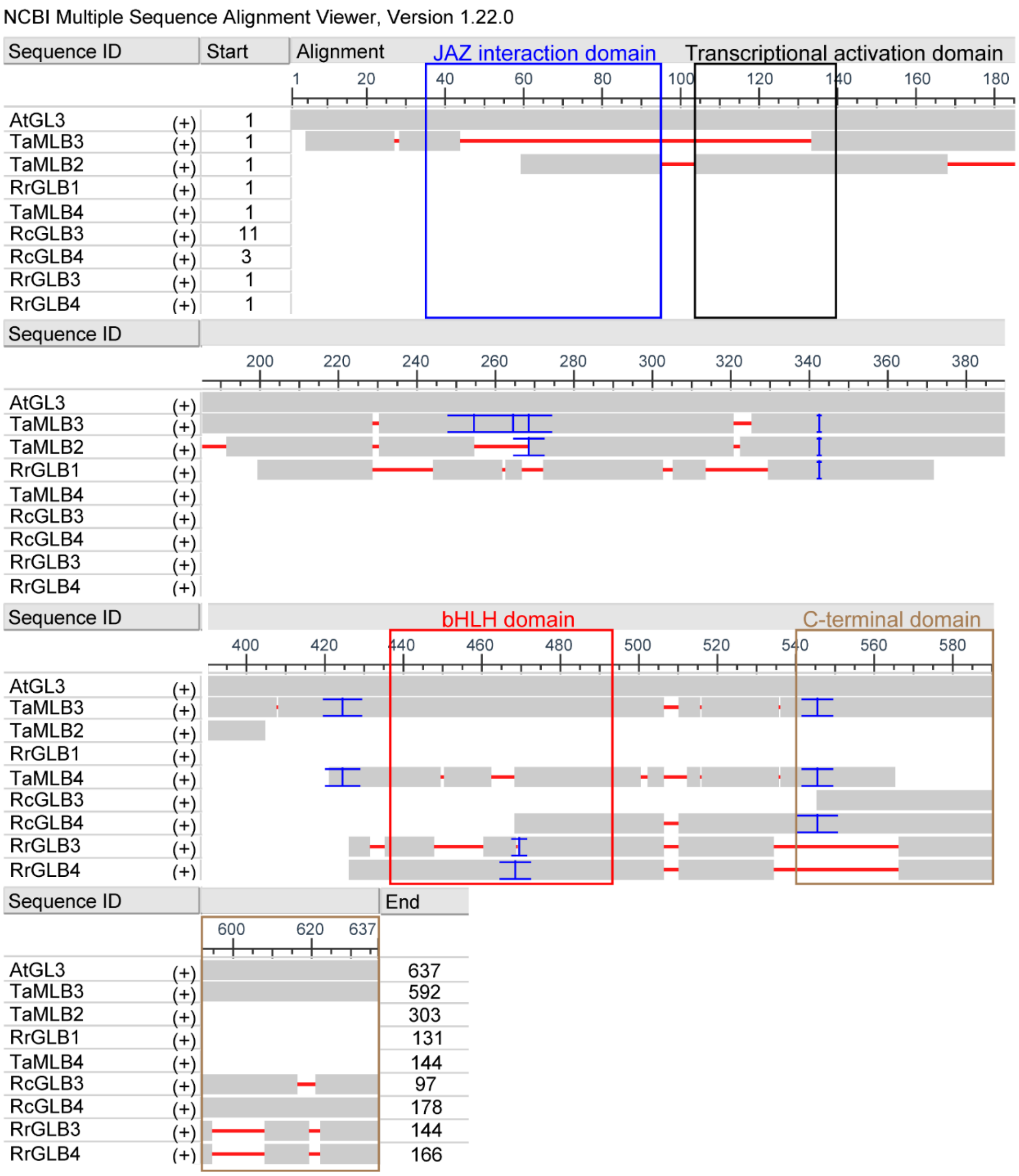


**Figure S12**

Sequence alignment of truncated GLBs and MLBs. Truncated GLBs and MLBs are aligned to AtGL3 and visualized by NCBI multiple sequence alignment viewer. The JID, TAD, bHLH domain and C-terminal domain are highlighted.

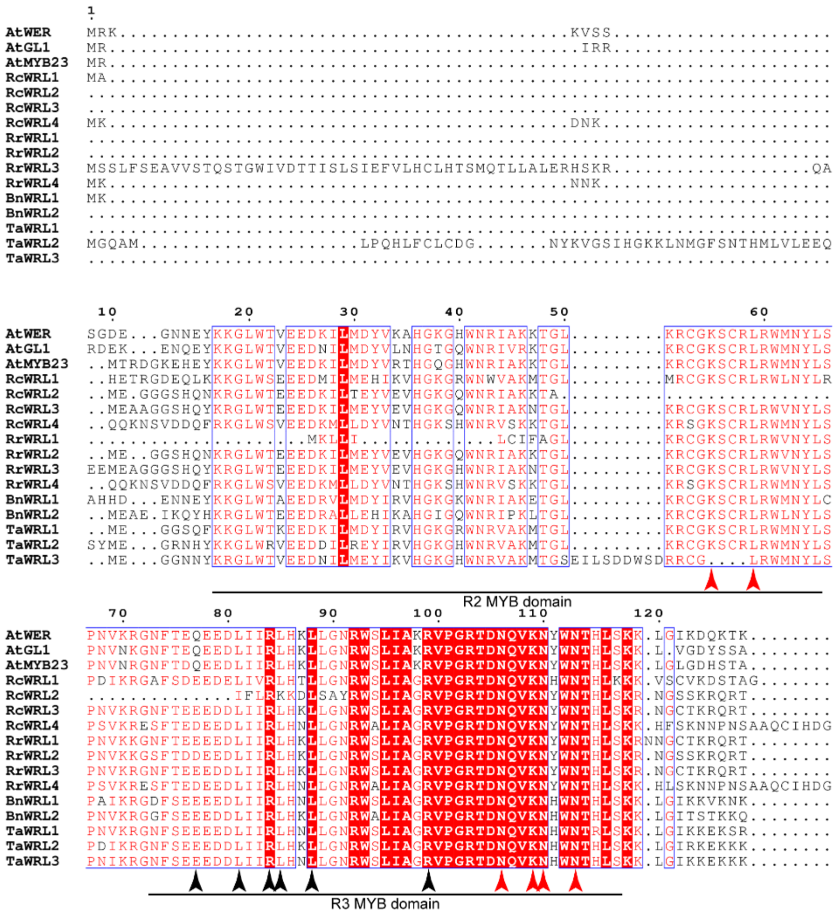

R2 MYB domain
R3 MYB domain

```
                            130                                            140
AtWER    ...................QSNG.....................................DIVYQINLP..
AtGL1    ..................VKTTG.....EDDD....SPPSL...................FITAATPSSCH
AtMYB23  ..................VKAAC......GVE....SPPSM...................ALITTTSSS..
RcWRL1   ..........FIKVQVSSCNANSFETVTPYKE....ENKSL...................LLSTQNMNC..
RcWRL2   ...................ESY.....APNPD....TRSSA.........TTLAPLSSNSNTLTTMENA..
RcWRL3   ...................QSY.....TPNPE....TNSET...................IFVTAAVNA..
RcWRL4   KR...CKDQNHFEDQVISSEFSS..NTTCRTK....ASSAV...................PLVTANKEV..
RrWRL1   ...................TAA.....MSTSSLVSLQPSKT...................II...PVNP..
RrWRL2   ...................ESY.....APNPD....TRSSACSKDSEAATTTLAPLSSNSNTLTTTENA..
RrWRL3   ...................QSYI....IPNPE....TNSET...................ILVTTVENA..
RrWRL4   KLKSCCKDQNHIEDQVISSEFSS..NTTCGTK....ASSAL...................PLVTANKEV..
BnWRL1   .................EMRVNS....QPLSKDQ..QEKDY...................FIPSGASLS..
BnWRL2   ......................................SRK...................PIDSHNTIAMT
TaWRL1   ..................VGSSS....LTHSREVG.ETQRL...................LIDSNSKLP..
TaWRL2   ..................VGASS....QAHSREVG.KTLSP...................TIDSNSKLP..
TaWRL3   ..................VGVSS....QTHSKEVG.LTLSP...................PIDSNSKLP..

                 150                 160          170
AtWER    .NPTETSEETKIS..NIVDN........NNI...LGDEIQEDHQG..SNYL....................
AtGL1    HQQENIYENIAKSFNGVVSA.SYEDKPKQEL...AQKDVLMATTNDPSHYY...GN...............
AtMYB23  HQEISGGKNSTLRFDTLVDE.S.KLKPKSKLVHATPTDVEVAATV..PNLF....................
RcWRL1   .......................QEATMCTQ...EAVDGEEELRS.....................F....
RcWRL2   EQETTVA....................DTSI....AQSADHAVVADGQKCG....S...............
RcWRL3   EQENTAVGNN...........NNNATGAASF....FQNAD.CFLANGA.......S...............
RcWRL4   .EETDTGQQSSQLG.......ALQANQPGEF....AQGETAAAD...DHG.....................
RrWRL1   EQETSAMENT...........TIISNALSS......HNAHACLVEDEQNYC....S...............
RrWRL2   EQET.......................NTSI....ARSAD.AVVADGQKCG....S...............
RrWRL3   EHENKNTAVG...........NNNATGAASF....SQNAD.PFLADGASS.....................
RrWRL4   .EETDTGQPSTQLG.......ALQANQPGEF....AQGETAAADEDDHDHG....................
BnWRL1   .EQPNTGNYTGDQAGIIGDKVTRDIGPKSSL.....EDVMMMSDNNDDNYE....................
BnWRL2   PVEEAGGQNCNVIMGRALEN.N.NIVVANDQGKPEEFEVPLECARN.EIFG...................P
TaWRL1   .CDNNGGSIGP..........KLNEGPQNTI...EARDTQ.ETTS..VCY.....................
TaWRL2   .NDSNGGD...............QEGPQNAI...GASDTQKQMMS..QCYGWTSGSQALCQAEGLGFDFS.
TaWRL3   .NDSNGGE................KGPQNAI...EASNTQEHMMN..ECFL....................

            180                   190                             200
AtWER    .SSLW........VHEDEF....ELSTLT.......N.MMDFID..............G...HCF......
AtGL1    .NALW........VHDDDF....ELSSLV.........MMNFAS..............GDVEYCL......
AtMYB23  .DTFW........VLEDDF....ELSSLT.........MMDFTN..............G...YCL......
RcWRL1   .SWMW.........................TNDPA.SPMELTLN..............G...SSDSWDW..
RcWRL2   .SLSW..........REQL..DEQMSSYQ.......SGLVQFSD..............E..FLQFDFGW..
RcWRL3   .SPVW..........RDQL..EQQMSIYQ.......S.LMQVSD..............E..FLQFDFGW..
RcWRL4   ....................................SCFMQYLD..............D...YTLHFVL..
RrWRL1   .SPTW..........R......EHMSSYQ.......N.LMQFSD..............E.....FDFGW..
RrWRL2   .SLSW..........REQL..DEQMSSYQ.......SGLVQFSD..............E..FLQFDFGW..
RrWRL3   .......................................................................
RrWRL4   ....................................SCFMQYLD..............D...YTLHFVL..
BnWRL1   .SSFW.......ITSADDFLINPNYSSYY.......D.VLEPLE..............E...QCLDFVW..
BnWRL2   ESPLM.........MRDDR....DFSDIILSRNPGNS.IVEFLD..............G...FCIPQLD..
TaWRL1   .ESLE........LLDDNF....NLNSPN.........LMELLD..............G...YPLELVW..
TaWRL2   .LPLWGQPRTGFPDFFD.V....ALRRRT.........LSNIHEIRSFSWQQSYMEGRN...HYKKGLWRV
TaWRL3   .SPLW........DFTDNL....NLNTPC.........LMELFD..............G...YPLDAVW..
```

```
AtWER     .......................................................................
AtGL1     .......................................................................
AtMYB23   .......................................................................
RcWRL1    ..............................................................FGH...DST
RcWRL2    ..............................................................NNI...MSS
RcWRL3    ..............................................................DN....MSS
RcWRL4    ..............................................................NNN..KRN.
RrWRL1    ..............................................................SN....MNS
RrWRL2    ..............................................................NN....MSS
RrWRL3    .......................................................................
RrWRL4    ..............................................................NNN..KRNA
BnWRL1    ........................................................D.....IGL......
BnWRL2    ........................................................D.....HQM...YWH
TaWRL1    ..............................................................HGL......
TaWRL2    EEDDILREYIRVHGKGHWNRVAKMTGLKRCGKSCRLRWMNYLSPDIKRGNFSEEEDDLIIRLHKLLGNRWS
TaWRL3    ..............................................................HGL......


AtWER     .......................................................................
AtGL1     .......................................................................
AtMYB23   .......................................................................
RcWRL1    IYDDL..................................................................
RcWRL2    LFSD...................................................................
RcWRL3    LFSD...................................................................
RcWRL4    AAEGM..................................................................
RrWRL1    LFSG...................................................................
RrWRL2    LFSD...................................................................
RrWRL3    .......................................................................
RrWRL4    AAEGM..................................................................
BnWRL1    .......................................................................
BnWRL2    YF.....................................................................
TaWRL1    .......................................................................
TaWRL2    LIAGRVPGRTDNQVKNHWNTHLSKKLGIRKEKKKVGASSQAHSREVGKTLSPTIDSNSKLPNDSNGTATIS
TaWRL3    .......................................................................


AtWER     ..................
AtGL1     ..................
AtMYB23   ..................
RcWRL1    ..................
RcWRL2    ..................
RcWRL3    ..................
RcWRL4    ..................
RrWRL1    ..................
RrWRL2    ..................
RrWRL3    ..................
RrWRL4    ..................
BnWRL1    ..................
BnWRL2    ..................
TaWRL1    ..................
TaWRL2    NNASDDQSKLKLEAKADG
TaWRL3    ..................
```

**Figure S13**

Sequence alignment of WRLs. Conserved amino acids among some of the WRLs (>70%) are colored as red. Conserved amino acids among all WRLs are colored as white and shaded with red. Conserved regions shared by WRLs (>70%) are labelled by blue squares. The R2 and R3 MYB domains are highlighted. The black arrowheads point to the residues mediating interaction with bHLH proteins. The red arrowheads point to the residues binding to DNA.

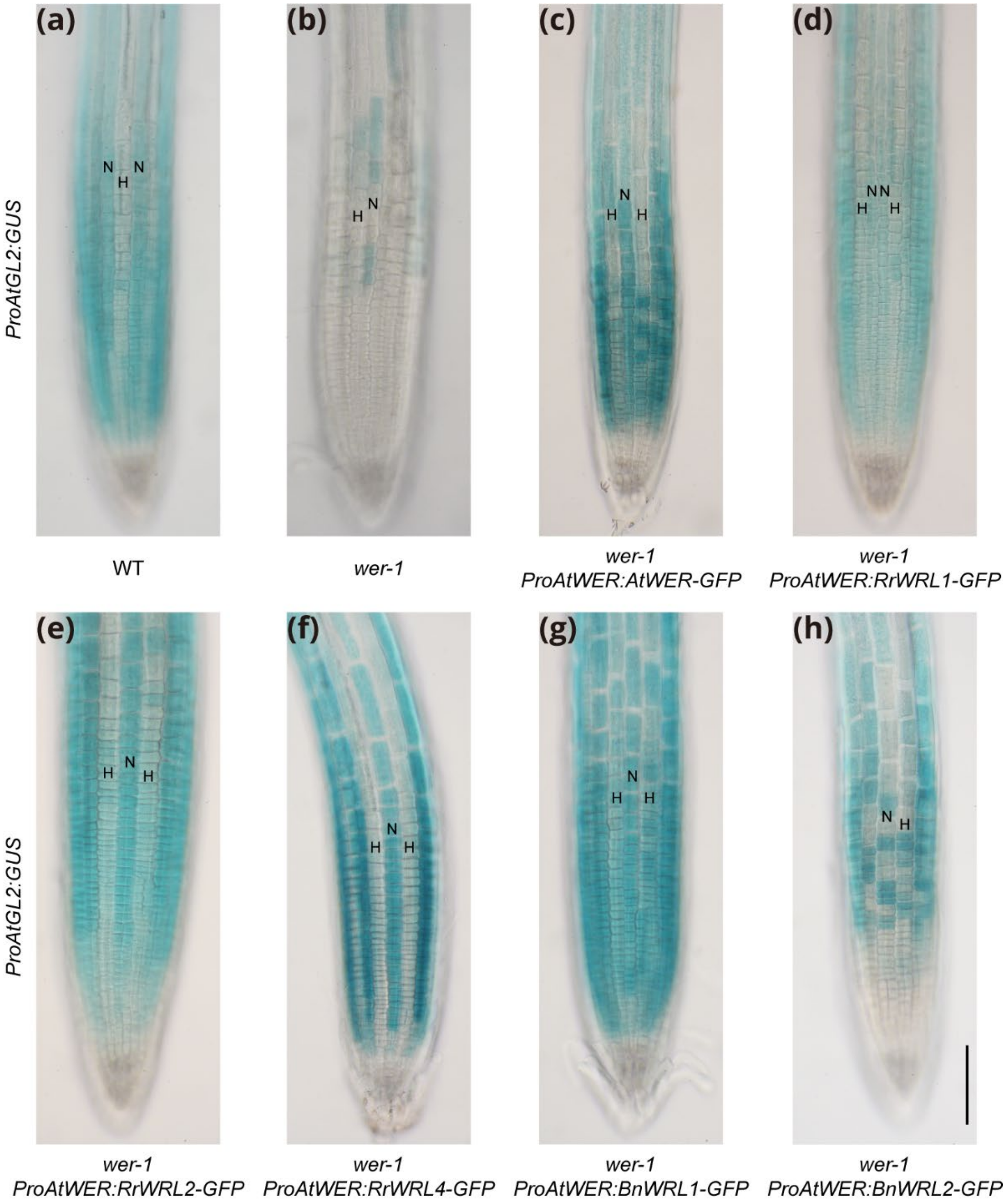


**Figure S14**

Expression of *ProAtGL2:GUS* transcriptional reporter in Arabidopsis lines bearing *ProAtWER:WRL-GFP* constructs. N-position cell files are labelled by N, and H-position cell files are labelled by H. All transgenic lines shown are representative T3 individuals. Bar=100um.

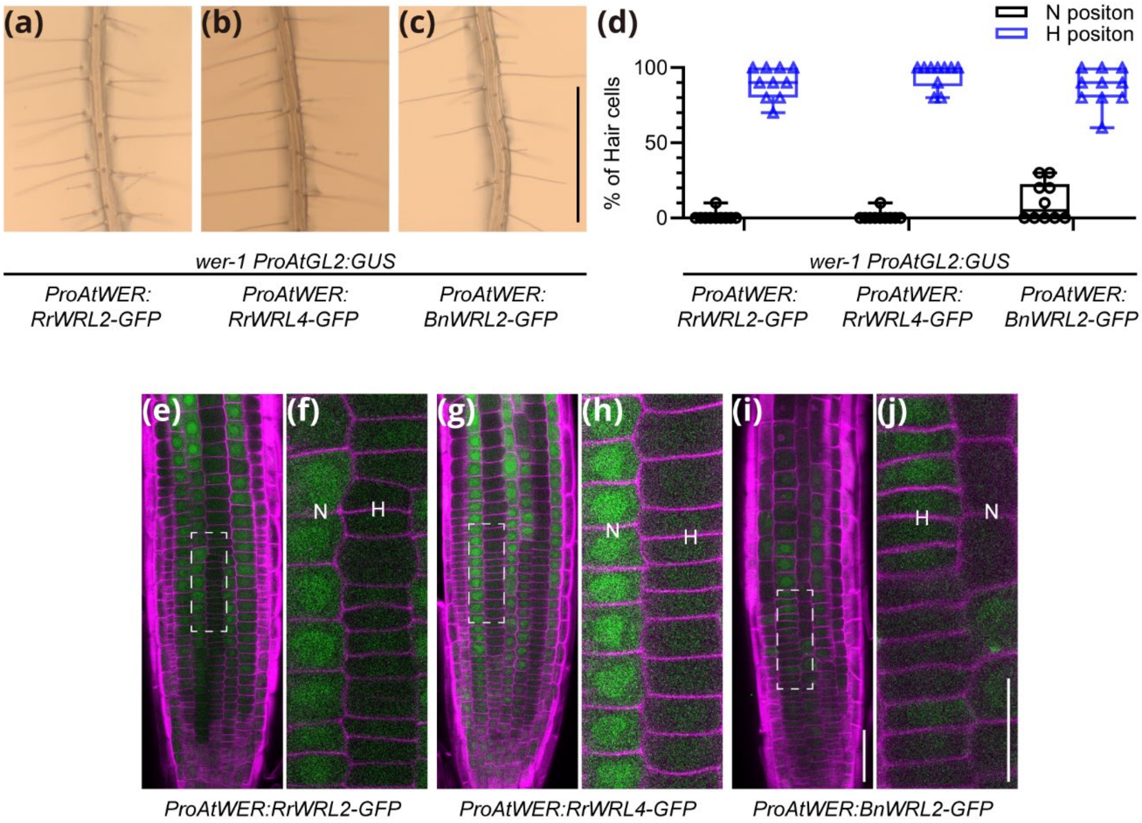


**Figure S15**

Functional analysis of additional *WRL*s. (**a**-**c**) Root hair phenotypes of Arabidopsis *wer-1* mutants bearing *ProAtWER:WRL-GFP* constructs. Each line also bears a *ProAtGL2:GUS* transgene. All transgenic lines shown are representative T3 individuals. Bar=1mm. (**d**) Quantification of root epidermal cell phenotypes of the lines shown in (**a**-**c**). Root hair counting raw data and information of other independent lines are listed in **Dataset 1**. (**e**, **g**, **i**) Confocal images of the lines shown in (**a**-**c**). (**f**, **h**, **j**) Zoom in view of the regions highlighted by dash lines in (**e**, **g**, **i**). N-position cell files are labelled by N, and H-position cell files are labelled by H. Bar=25um. GFP signals are pseudocolored as green, CalcoFluor White signals are pseudocolored as magenta.

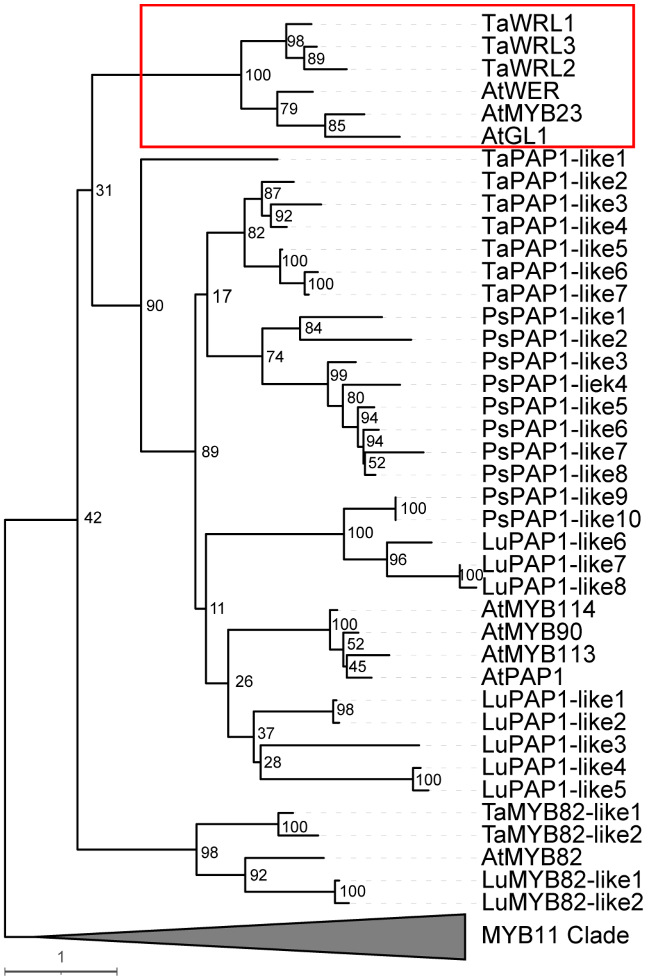


**Figure S16**

Absence of WRLs from Type I species *L.usitatissimum* and *P.sativum*. Maximum likelihood tree of WRLs, AtPAP1-like proteins and AtMYB82-like proteins. The tree was rooted to the MYB11 clade. The scale bar refers to amino acid substitutions per site. The WRL clade is highlighted. *A.thaliana*, *T.aralioide*, *L.usitatissimum*, and *P.sativum* proteins were used for tree construction. Lu, *L.usitatissimum* proteins; Ps, *P.sativum* proteins.

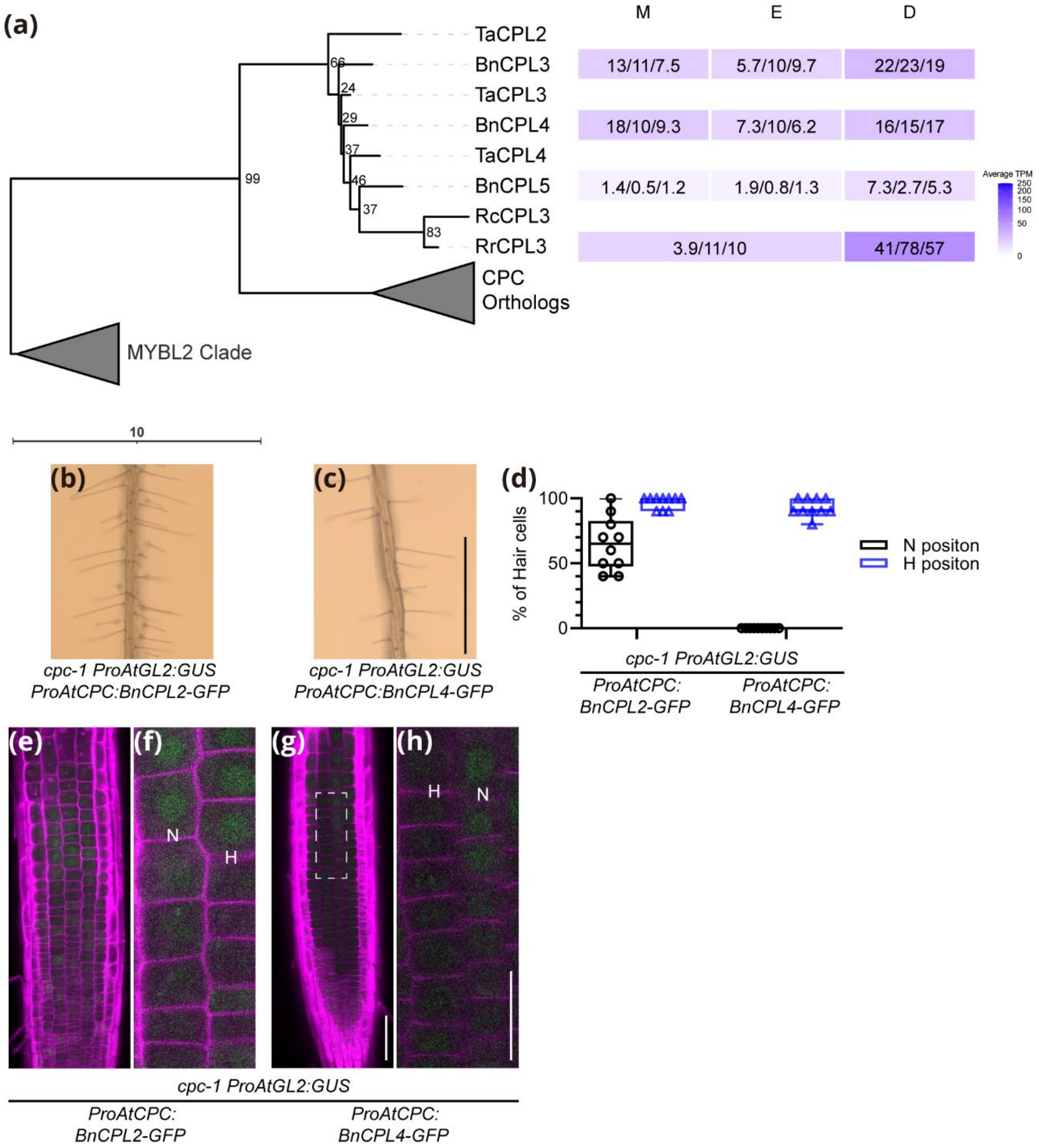


### Figure S17

Identification and functional analysis of additional *CPL*s. (**a**) Left, maximum likelihood tree of CPLs. The tree was rooted the MYBL2 clade. The scale bar refers to amino acid substitutions per site. The CPC ortholog clade shown in Fig. 5 was collapsed. Right, Gene expression levels in root developmental zones. TPM values of 3 RNA-seq replicates are shown in each box, and the average values are visualized by heatmaps after square root transformation. (**b**) Root hair phenotypes of *cpc-1* mutant seedlings bearing *ProAtCPC:BnCPL2-GFP* constructs. A representative T3 individual is shown. Bar=1mm. (**c**) Root hair phenotypes of *cpc-1* mutant seedlings bearing *ProAtCPC:BnCPL4-GFP* constructs. A representative T3 individual is shown. Bar=1mm. (**d**) Quantification of root epidermal cell phenotypes of the lines shown in (**b**) and (**c**). Root hair counting raw data and information of other independent lines are listed in

**Dataset 1**. (**e**, **g**) Confocal images of the lines shown in (**b**) and (**c**). Bar=50um. (**f**, **h**) Zoom in view of the regions highlighted by dash lines in (**e**, **g**). Bar=25um.

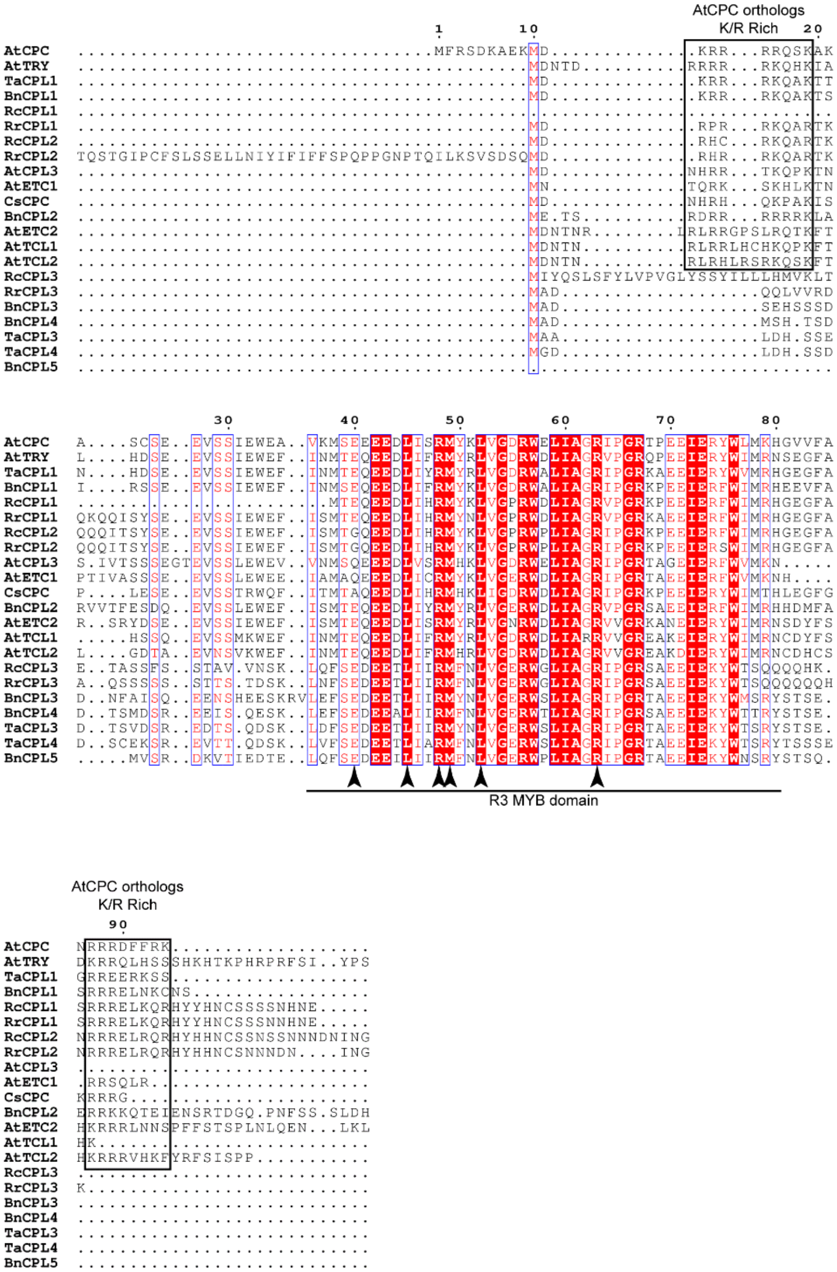


**Figure S18**

Sequence alignment of CPLs. TaCPL2 was not included due to extended C-terminal sequences. Conserved amino acids among some of the CPLs (>70%) are colored as red. Conserved amino acids among all CPLs are colored as white and shaded with red. Conserved regions shared by CPLs (>70%) are labelled by blue squares. The R3 MYB domain and the N- and C-terminal K/R rich regions found in AtCPC othologs are highlighted. The arrowheads point to the residues mediating interaction with bHLH proteins.

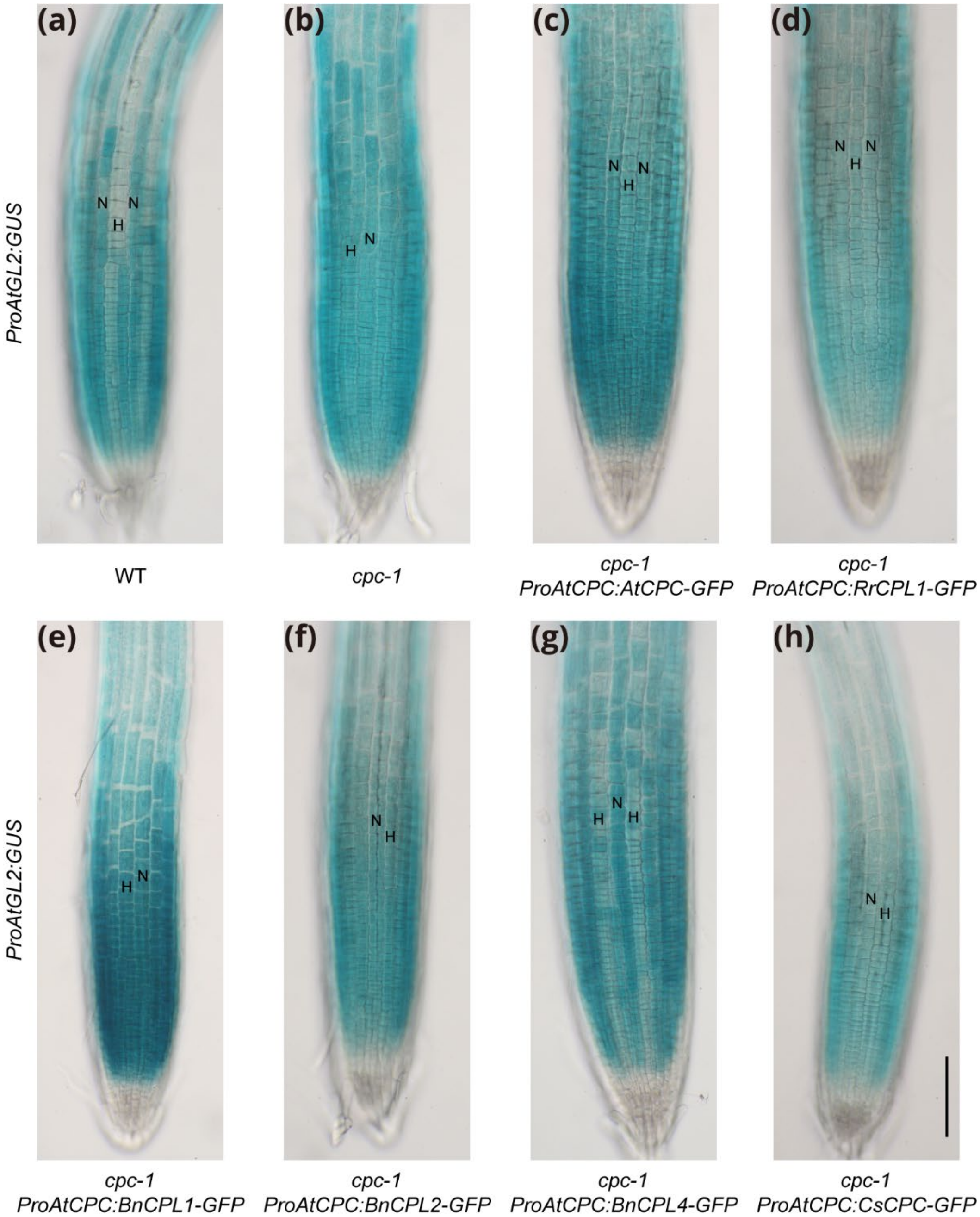


**Figure S19**

Expression of *ProAtGL2:GUS* transcriptional reporter in Arabidopsis lines bearing *ProAtCPC:CPL-GFP* constructs. N-position cell files are labelled by N, and H-position cell files are labelled by H. Bar=100um.

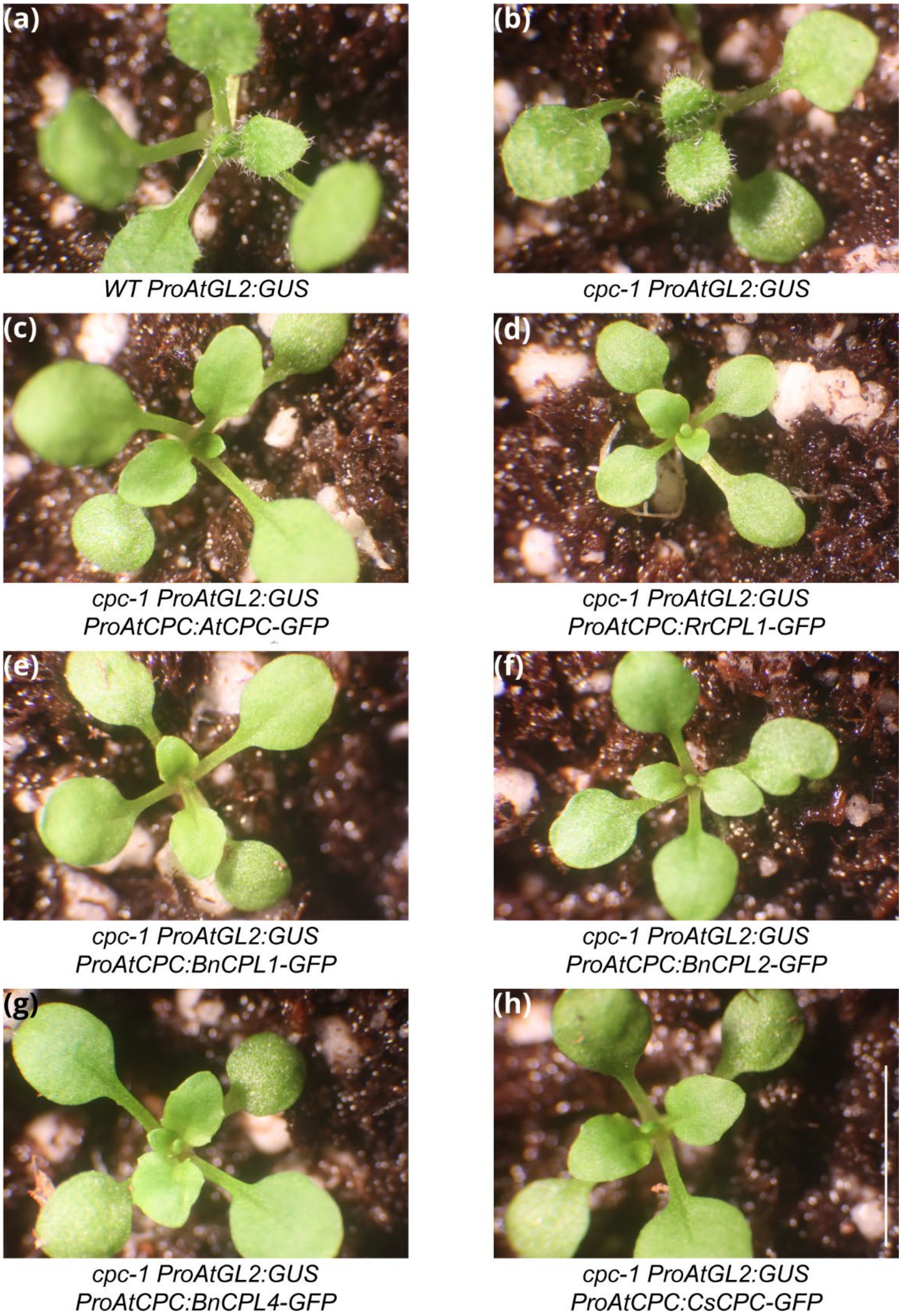


**Figure S20**

Trichome phenotypes of lines bearing *ProAtCPC:CPL-GFP* constructs. All transgenic lines shown are representative T3 individuals. Bar=5mm.

**Methods S1** Root hair counting assay

Seedling roots of the plants were stained with a dye containing 1% toluidine blue and 1% Triton X-100 briefly. The roots were then mounted in water and observed under a bright-field microscope (Laborlux S; Leica). For each individual root, two regions were counted: Region 1, which begins at the position where root hairs start to emerge, and Region 2, which is a randomly selected area of the mature root where all root hairs are fully grown and proximal to hypocotyl. For each region, 5 continuous N-position cells and neighboring H-position cells were analyzed for presence of root hairs.

**Methods S2** Plastic embedding and transverse sectioning of plant roots

Seedling roots were first fixed overnight at 4°C with 3% glutaraldehyde and 50mM NaPO4 (pH 6.8). The seedlings were then washed with water and embedded in 2.5% agarose. The agarose blocks were dehydrated in a graded ethanol series (10%, 20%, 50%, 70%, 90%, 100%, 100%, 15 min each). The blocks were then treated with propylene oxide for 10 min in a glass vial with closed lid, followed by infiltration with EMbed 812 (Electron Microscopy Sciences) mix/propylene oxide (1:1) for 1 h with the lid open. The infiltration was step repeated with EMbed 812 mix/propylene oxide (1:1) for 1 h with the lid closed. The blocks were then transferred into a new vial and infiltrated with pure Embed 812 mix overnight. The blocks were trimmed and orientated in plastic molds filled with pure Embed 812 mix and polymerized at 60°C for 16 h. The Embed 812 mix was prepared as follows: 11.0 g Embed 812, 7.275 g NMA and 4.52 g DDSA were mixed well, then 0.35 g DMP-30 was added and mixed thoroughly. Transverse sections (0.95 um in thickness) were cut using glass knives with an ultramicrotome (Ultracut E; Leica-Reichert). The sections were stained with 1% toluidine in 1% sodium borate solution then observed and photographed under a bright-field microscope.

**Methods S3** Generation of plasmids

The *pCAM-EYFP-C1* plasmids were digested with EcoRI and BglII to remove the *CaMV35S* promoter and *EYFP* cDNA sequences (CDS). Synthesized nucleotides were ligated into the plasmids to generate an EcoRI-SalI-PstI-BamHI-MluI-BglII multiple cloning site. The resulting vector was named *pCAM-YZ1*. The *bar* gene, encoding phosphinothricin acetyltransferase, was cloned from binary vector *pCB302* (Xiang *et al.*, 1999) and inserted into *pCAM-YZ1* to replace the *HygR* gene. The resulting vector was named *pCAM-YZ2*.

To construct *ProAtRHD6:GFP-RLB* plasmids, a 2.9 kb genomic fragment upstream of *AtRHD6* open reading frame (ORF) was cloned as *ProAtRHD6*. A 1 kb genomic fragment downstream of *AtRHD6* ORF was cloned as *AtRHD6* 3' untranslated region (UTR). *mGFP5*, which encodes a green fluorescent protein was cloned with MluI and SacI restriction enzyme sites added to the 3' end. *pCAM-YZ1* was digested with SalI and BglII and all mentioned fragments were incorporated into the vector to generate a *ProAtRHD6:GFP* vector backbone, using the NEBuilder HiFi DNA Assembly Cloning kit (New England Biolabs). The *RLB* CDSs were cloned from cDNA libraries. *AtRHD6* and *BnRLB* CDSs were ligated into the *ProAtRHD6:GFP* backbone using the MluI and SacI sites. The *RrRLB1* CDS was incorporated into the backbone using the NEBuilder HiFi DNA Assembly Cloning kit.

To construct *ProAtGL2:GFP-GLH* plasmids, a 2.4 kb genomic fragment upstream of *AtGL2* ORF was cloned as *ProAtGL2*. A 459 bp genomic fragment downstream of *AtGL2* ORF was cloned as *AtGL2 3'UTR*. *mGFP5* was cloned with BglII and PstI restriction enzyme sites added to the 3' end. *pCAM-YZ1* was digested with EcoRI and BglII and all mentioned fragments were incorporated into the vector to generate a *ProAtGL2:GFP* vector backbone, using the NEBuilder HiFi DNA Assembly Cloning kit. The *GLH* CDSs were then cloned from cDNA libraries. *AtGL2*, *BnGLH*, and *RrGLH1* CDSs were incorporated into the *ProAtGL2:GFP* backbone using the NEBuilder HiFi DNA Assembly Cloning kit. *CsGLH1* CDS was ligated into the backbone using BglII and NsiI sites.

To construct *ProAtWER:WRL-GFP* plasmids, a 4 kb genomic fragment upstream of *AtWER* ORF was cloned as *ProAtWER*. A 1.1 kb genomic fragment downstream of *AtWER* ORF was cloned as *AtWER 3'UTR*. *BnWRL1* CDS was first cloned with a PstI site added to 5' end. *mGFP5* was cloned with a BamHI site added to 5' end. *pCAM-YZ1* was digested with SalI and BglII and all mentioned fragments were incorporated into the plasmid to generate a *ProAtWER:BnWEL1-GFP* vector, using the NEBuilder HiFi DNA Assembly Cloning kit. The *BnWRL1* CDS was then cut from the vector using PstI and

BamHI. Other *WRL* CDSs were then cloned from cDNA libraries. *AtWER*, *RrWRL2, and RrWRL4* CDSs were ligated into the plasmid using PstI and BamHI sites. *BnWRL2* and *RrWRL1* CDSs were ligated into the plasmid using NsiI and BamHI sites.

To construct *ProAtCPC:CPL-GFP* plasmids, a 2 kb genomic fragment upstream of *AtCPC* ORF was cloned as *ProAtCPC*. A 1 kb genomic fragment downstream of *AtCPC* ORF was cloned as *AtCPC 3'UTR*. *mGFP5* was cloned with a MluI restriction enzyme site added to the 5' end and a BsrGI site added to 3'end. *pCAM-YZ2* was digested with EcoRI and BglII and all mentioned fragments were incorporated into the vector to generate a *ProAtCPC:GFP* vector backbone, using the NEBuilder HiFi DNA Assembly Cloning kit. The backbone was digested with MluI and treated with calf intestinal alkaline phosphatase (CIP). The *CPL* CDSs were cloned from cDNA libraries. *AtCPC* CDS was incorporated into the *ProAtCPC:GFP* backbone using the NEBuilder HiFi DNA Assembly Cloning kit. All other *CPL* CDS was ligated into the backbone using flanking MluI sites.

To construct *Promoter:*$Sp^{WAK2}$*-mCherry-HDEL* (Nelson *et al.*, 2007) plasmids, the sequence encoding the signal peptide of *AtWAK2 (*$Sp^{WAK2}$) which localizes the protein to ER was cloned from an Arabidopsis cDNA library. *mCherry* which encodes a red fluorescent protein was cloned with sequences encoding an ER retention signal HDEL added to 3' end. *ProAtWER:AtWER-GFP* plasmid was digested with PstI and MluI to remove the *AtWER-GFP* CDS. The $Sp^{WAK2}$ and *mCherry-HDEL* fragments were incorporated into the digested vector using the NEBuilder HiFi DNA Assembly Cloning kit to generate the *ProAtWER:*$Sp^{WAK2}$*-mCherry-HDEL* construct. The *ProAtCPC:BnCPL1-GFP* vector was digested with MluI and BsrGI to remove the *BnCPL1-GFP* CDS. The cloned *SpWAK2-mCherry-HDEL* cassette was incorporated into the digested vector to generate the *ProAtCPC:*$Sp^{WAK2}$*-mCherry-HDEL* construct using NEBuilder HiFi DNA Assembly Cloning kit. The *SpWAK2-mCherry-HDEL* cassette was cloned with a SalI site added to 5' end and a MluI site added to 3' end, and ligated into *pCAM-YZ1* using these sites. A 3 kb genomic fragment upstream of *BnGLH* ORF was cloned as *ProBnGLH*. A 3 kb genomic fragment upstream of *BnWRL1* ORF was cloned as *ProBnWRL1*. A 2 kb genomic fragment starting from 65 bp upstream of *BnWRL2* ORF was cloned as *ProBnWRL2*. A 2 kb genomic fragment upstream of

*BnCPL1* ORF was cloned as *ProBnCPL1*. All the promoter sequences were incorporated into the backbone upstream of *SpWAK2-mCherry-HDEL* cassette using the NEBuilder HiFi DNA Assembly Cloning kit.

**Methods S4** RNA Isolation and sequencing

*B.nivea*, *A.thaliana* and *C.sativus* RNA was isolated using Qiagen RNeasy Plant Mini Kit, and *R.rosea* zonation RNA was isolated using Takara NucleoSpin RNA Plant and Fungi kit. On-column DNA digestion was performed using Qiagen RNase-Free DNase Set or Takara rDNase Set according to the manufacturer's instructions. The cDNA libraries used for gene cloning were synthesized using Invitrogen SuperScript First-Strand synthesis system.

Takara SMART-Seq v4 Ultra Low Input RNA Kit was used for *R.rosea* RNA library construction. Illumina TruSeq Kit was used for *B.nivea* RNA library construction. The *B.nivea* RNA libraries were sequenced using Illumina HiSeq2500 and HiSeq4000 system (50 Cycles, 50 bp Single End) and *R.rosea* RNA libraries were sequenced using Illumina NovaSeq 6000 system (S4 flow cell, 300 cycles, 150 bp Paired End).

**Methods S5** RNA-seq analysis

All reads were first analyzed using FastQC (Version 0.11.5) (https://www.bioinformatics.babraham.ac.uk/projects/fastqc/). The *A.thaliana*, *C.sativus*, and *B.nivea* reads were trimmed using TrimGalore (Version 0.6.7) (https://www.bioinformatics.babraham.ac.uk/projects/trim_galore/). The 5' end 15 bp of the reads were trimmed as previously described (Huang & Schiefelbein, 2015). These processed reads were then used for downstream analysis.

Ribosomal RNA (rRNA) was detected after initial quality check of *R.rosea* raw reads and these rRNA reads were removed by mapping raw reads to rRNA database (Quast *et al.*, 2013) (SILVA 138.1, LSU_Parc and SSU_Parc) with Bowtie2 (Version 2.4.2) (Langmead & Salzberg, 2012). Erroneous Kmers were removed by rcorrector (Version 1.0.4) (Song & Florea, 2015). The corrected reads were trimmed by TrimGalore (Version 0.6.7) (https://www.bioinformatics.babraham.ac.uk/projects/trim_galore/). *De novo* assembly was performed with Trinity (Version 2.12.0, default mode), Trinity genome guided mode (reads first mapped to *R.crenulata* genome using Bowtie2) (Grabherr *et al.*, 2011), and rnaSPAdes (Version 3.15.5) (Bushmanova *et al.*, 2019). The three resulting transcriptomes were concatenated using EvidentialGene (Gilbert, 2019) to generate the final assembly. Quality of the final assembly was assessed using BUSCO (Version 4.0.6) (Manni *et al.*, 2021). The processed reads were mapped back to the assembly using Bowtie2 (Version 2.4.2) (Langmead & Salzberg, 2012) followed by quantification using RSEM (Version 1.3.3) (Li & Dewey, 2011).

The *R.rosea* transcriptome was annotated using Trinotate (Version 3.2.2) (Bryant *et al.*, 2017). In brief, TransDecoder (Version 5.5.0) (https://github.com/TransDecoder/TransDecoder) was used to predict peptides from the assembly, SignalP 6.0 (Teufel *et al.*, 2022) was used to predict signal peptides, and Tmhmm (Version 2.0) (Krogh *et al.*, 2001) was used to predict transmembrane domains. Homologies of the assembled transcripts and predicted peptides were identified by searching the Uniprot database using NCBI BLAST+ (Version 2.90) (Camacho *et al.*, 2009) . HMMER (Version 3.3.2) (http://hmmer.org/) was used to identify protein domains. All these results were loaded into Trinotate SQLite Database to generate the annotation report.

For all species, read counts quantified by RSEM were imported into R using tximport (Version 1.26.0) and normalized as previously described (Soneson *et al.*, 2015). Differential expression analysis was performed using edgeR (Version 3.40.0) (Robinson *et al.*, 2010). In brief, dispersions were estimated using the GLM method (estimateDisp() ), and differential expression was determined using quasi-likelihood F-tests (glmQLFit() and glmQLFTest() ). Genes with a $|\log2(\text{fold-change})| \geq 1$ and a false discovery rate $\leq 0.05$ between the compared developmental zones were retained and present in **Dataset 4**.

**Methods S6** Phylogenetic analysis

The annotations of *A.thaliana*, *B.nivea*, *R.crenulata*, *C.sativus*, *T.aralioides*, *L.usitatissimum and P.sativum* genomes and *R.rosea* transcriptome were used for phylogenetic analysis (Lamesch *et al.*, 2012; Wang *et al.*, 2012; Fu *et al.*, 2017; Torrens-Spence *et al.*, 2018; Kreplak *et al.*, 2019; Li *et al.*, 2019; Strijk *et al.*, 2019; Wang *et al.*,

2021). The pipeline was modified from one previously described (Huang & Schiefelbein, 2015) . In brief, NCBI BLAST+ (Version 2.90) (Camacho *et al.*, 2009) was used to query the protein of interest against predicted proteins of all selected species. The outputs with e-value ≤ 1 were clustered by SWIPE (Version 2.1.0) (swipe -M BLOSUM62 -m 8 -e 1 -v 1000 -b 1000) (Rognes, 2011) and the longest protein of each gene model was retained. All the sequences were aligned by MAFFT (Version 7.407) (mafft --ep 0 --genafpair --maxiterate 1000 --namelength 100 --reorder) (Katoh & Standley, 2013). FastTree (Version 2.1.11) (fasttree -gamma) was used to infer an approximately-maximum-likelihood tree (Price *et al.*, 2009). Sequences in a well-supported clade containing the protein of interest, together with proteins in the sister clade or an outgroup were selected and aligned again using MAFFT. The best substitutional model was determined by ModelFinder integrated in IQ-tree (Version 2.1.2) (Kalyaanamoorthy *et al.*, 2017). The maximum likelihood tree was then inferred with raxml-ng (Version 1.1.0) (raxml-ng --model best_model --all --bs-trees autoMRE) (Kozlov *et al.*, 2019).

The heat maps aligned to the phylogenetic tress were generated using ggplot2 (Wickham, 2009). TPMs of all tested genes from examined species were normalized together for color gradient.

The phylogenetic trees together with heat maps were visualized with iTOL (Letunic & Bork, 2021), FigTree (http://tree.bio.ed.ac.uk/software/figtree/), and Dendroscope (Huson & Scornavacca, 2012) and modified in Adobe Illustrator and Acrobat.

The sequence alignments were visualized using ESPript3 (https://espript.ibcp.fr) (Robert & Gouet, 2014).
